# An Alanine Aminotransferase is Required for Polysaccharide Regulation and Resistance of *Aspergillus fumigatus* Biofilms to Echinocandin Treatment

**DOI:** 10.1101/2021.06.03.446912

**Authors:** Joshua D. Kerkaert, François Le Mauff, Benjamin R. Wucher, Sarah R. Beattie, Elisa M. Vesely, Donald C Sheppard, Carey D. Nadell, Robert A. Cramer

## Abstract

Alanine metabolism has been suggested as an adaptation strategy to oxygen limitation in organisms ranging from plants to mammals. Within the pulmonary infection microenvironment *A. fumigatus* forms biofilms with steep oxygen gradients defined by regions of oxygen limitation. A significant increase in alanine levels was observed in *A. fumigatus* cultured under oxygen limiting conditions. An alanine aminotransferase, AlaA, was observed to function in alanine catabolism and is required for several aspects of *A. fumigatus* biofilm physiology. Loss of *alaA*, or its catalytic activity, results in decreased adherence of biofilms through a defect in the maturation of the extracellular matrix polysaccharide galactosaminogalactan (GAG). Additionally, exposure of cell wall polysaccharides is also impacted by loss of *alaA* and loss of AlaA catalytic activity confers increased biofilm susceptibility to echinocandin treatment which is correlated with enhanced fungicidal activity. The increase in echinocandin susceptibility is specific to biofilms and chemical inhibition of *alaA* by the alanine aminotransferase inhibitor β-chloro-L-alanine is sufficient to sensitize *A. fumigatus* biofilms to echinocandin treatment. Finally, loss of *alaA* increases susceptibility of *A. fumigatus* to *in vivo* echinocandin treatment in a murine model of invasive pulmonary aspergillosis. Our results provide insight into the interplay of metabolism, biofilm formation, and antifungal drug resistance in *A. fumigatus* and describes a mechanism of increasing susceptibility of *A. fumigatus* biofilms to the echinocandin class of antifungal drugs.

**eLife Digest:** *Aspergillus fumigatus* is a ubiquitous filamentous fungus that causes an array of diseases depending on the immune status of an individual, collectively termed aspergillosis. Antifungal therapy for invasive pulmonary aspergillosis (IPA) or chronic pulmonary aspergillosis (CPA) is limited and too often ineffective. This is in part due to *A. fumigatus* biofilm formation within the infection environment and the resulting emergent properties, particularly increased antifungal resistance. Thus, insights into biofilm formation and mechanisms driving increased antifungal drug resistance are critical for improving existing therapeutic strategies and development of novel antifungals. In this work, we describe an unexpected observation where alanine metabolism, via the alanine aminotransferase AlaA, is required for several aspects of *A. fumigatus* biofilm physiology including resistance of *A. fumigatus* biofilms to the echinocandin class of antifungal drugs. Importantly, we observed that chemical inhibition of alanine aminotransferases is sufficient to increase echinocandin susceptibility and that loss of *alaA* increases susceptibility to echinocandin treatment in a murine model of IPA.

## Introduction

*Aspergillus fumigatus* is a ubiquitous filamentous fungus with a prominent ecological role in the decomposition of organic carbon, that is easily isolated from compost piles and similar environments (Gugnani, 2003). Within compost piles a complex set of microenvironments can emerge along temperature and nutrient gradients that naturally form as saprophytes become metabolically active (Di Piazza et al., 2020; Sánchez et al., 2017). Thus, *A. fumigatus* has evolved a significant degree of metabolic flexibility and thermotolerance (Bhabhra & Askew, 2005; Ries et al., 2018). However, these saprophytic fitness traits also increase the fungus’ pathogenic potential leading to *A. fumigatus* being the causative agent of a variety of immune-status dependent human diseases (Casadevall, 2017; Kanj et al., 2018; Robert & Casadevall, 2009), with the most lethal disease manifestation being Invasive Pulmonary Aspergillosis (IPA).

IPA occurs primarily in individuals with a suppressed innate immune system, such as individuals undergoing solid-organ transplantation or chemotherapy (Kanj et al., 2018; Kousha et al., 2011) Tragically, antifungal therapy options for IPA remain limited and are often ineffective, with recent clinical trials reporting 12-week mortality rates of 28-45% depending on therapeutic regimen and host immune status (Herbrecht et al., 2015; Maertens et al., 2016, 2021; Marr et al., 2015). One class of antifungal drugs, the echinocandins, inhibits synthesis of cell wall β-glucans and are better tolerated by patients than drugs belonging to the azole or polyene classes of antifungals. In many pathogenic yeasts, such as *Candida albicans*, echinocandin treatment has fungicidal activity and is utilized as a first-line treatment. While echinocandins yield some level of cell lysis when applied to *A. fumigatus* hyphae, these drugs are primarily fungistatic against *A. fumigatus* as it is intrinsically tolerant to echinocandins and will exhibit residual growth even at high concentrations of drug (Moreno-Velásquez et al., 2017). In many cases this tolerance results in a paradoxical phenomenon where the fungus will recover growth as the concentration of drug increases beyond a minimal effective concentration (MEC) (Aruanno et al., 2019; Moreno-Velásquez et al., 2017; Wagener & Loiko, 2018). Thus, echinocandins have primarily been utilized as a salvage therapy for IPA and strategies to increase their efficacy in treatment of IPA are potentially of great clinical significance.

Recent studies have shown that *A. fumigatus* forms robust biofilms within the infection environment (Kowalski et al., 2019; Loussert et al., 2010). The hyphae within the *A. fumigatus* biofilm are coated with the extracellular matrix polysaccharide galactosaminogalactan (GAG), which is composed of a heterogenous mixture of galactose and N-acetylgalactosamine. GAG functions as a primary adherence factor for *A. fumigatus* biofilms, as well as an immunomodulatory compound (Fontaine et al., 2011; Gravelat et al., 2013; Lee et al., 2015; Speth et al., 2019). After synthesis GAG requires partial deacetylation via the Agd3 deacetylase to function in both capacities and strains lacking the ability to either produce GAG or deacetylate GAG are unable to adhere to surfaces (Bamford et al., 2020; Gressler et al., 2019; Lee et al., 2016). While some transcriptional regulatory machinery surrounding GAG biosynthesis and maturation has been described, mechanisms underlying biofilm formation and ECM regulation remain to be fully defined (Chen et al., 2020; Gravelat et al., 2013).

Insights into *A. fumigatus* biofilm formation are of great importance as these biofilms have been shown to display clinically relevant emergent properties, including increased resistance to antifungal drugs (Kowalski et al., 2020; Mowat et al., 2008; Seidler et al., 2008). A major factor contributing to increased drug resistance is the formation of oxygen limited, hypoxic, microenvironments within the biofilm (Kowalski et al., 2020, 2021). These same hypoxic microenvironments have been observed to exist in the infection environment and the ability to adapt to oxygen limitation is essential for disease progression and full virulence (Grahl et al., 2011; Willger et al., 2008). While some transcriptional regulators of *A. fumigatus* oxygen adaptation have been identified (Chung et al., 2014; Hagiwara et al., 2017; Willger et al., 2008), how the fungus metabolically adapts to low oxygen and how these metabolic pathways go on to impact broader *A. fumigatus* physiology remain to be fully appreciated. In order to examine metabolic pathways that are potentially important for low oxygen adaptation we conducted a metabolomics experiment looking at the impact of acute exposure to oxygen limitation.

Analysis of metabolomics data described in this work, combined with published transcriptomics data, suggests a role for alanine metabolism in low oxygen adaptation (Barker et al., 2012; Chung et al., 2014; Hillmann et al., 2014; Losada et al., 2014). Alanine metabolism has been associated with adaptation to oxygen limitation in numerous organisms ranging from plant roots adapting to waterlogging (Lothier et al., 2020; Rocha et al., 2010) to exercise induced oxygen deprivation in muscle cells (Felig, 1973). Importantly, alanine is also one of the handful of amino acids detectable in human bronchoalveolar lavage (BAL) fluid and bronchial wash (BW) fluid, indicating it is readily available in the airway environment (Surowiec et al., 2016) and may serve as a potential carbon or nitrogen source for *A. fumigatus*. Here, we explore the role of fungal alanine metabolism via the alanine aminotransferase AlaA in *A. fumigatus*. Alanine aminotransferases catalyze the interconversion of pyruvate and alanine utilizing glutamate as an amino-group donor and thus participates in both carbon and nitrogen metabolism. While we observe that AlaA-mediated metabolic reactions are not essential for low oxygen growth, AlaA catalytic activity is critical for normal biofilm physiology where oxygen gradients naturally form. Moreover, AlaA is critical for growth and full fitness when alanine is the sole carbon or nitrogen source. Unexpectedly*, alaA* is required for maturation of the ECM polysaccharide GAG and exposure of cell wall polysaccharides. Furthermore, deletion or inhibition of AlaA results in a striking reduction of echinocandin resistance in *A. fumigatus* biofilms both *in vitro* and *in vivo*.

## Results

### Acute exposure to low oxygen leads to the accumulation of carbohydrate metabolites and amino acids

To examine the metabolic impact of acute exposure to oxygen limitation, fungal batch cultures were grown for 24 hours in ambient oxygen followed by either continued incubation at ambient O_2_ or a 2-hour shift to an atmosphere of 0.2% O_2_. Intracellular metabolites were then extracted from biomass and relative quantities determined by LC-MS/MS. Acute exposure of fungal cultures to an oxygen limiting environment led to significant alterations in the relative abundance of 167 of 438 detected metabolites (38%) (Figure 1-figure supplement 1B, Supplemental File 1). Among these altered metabolites, an accumulation of several TCA cycle intermediates, lactate, and 4-amino-butyric acid (GABA) was observed. These metabolites are expected to increase during a rapid shift from a primarily oxidative metabolic state to a more fermentative metabolism (Figure 1A, Figure 1-figure supplement 1C). Other metabolites associated with fermentation such as acetate and ethanol were not detected via the method utilized, however other reports from similar culture conditions have observed ethanol fermentation as a major metabolic product during *A. fumigatus* growth in low oxygen conditions (Grahl et al., 2011). Additionally, decreased levels of glycolytic intermediates, in combination with increased intracellular trehalose content suggests a divergence of available carbon away from energy generation, in favor of carbon storage in the form of readily mobilized compounds, like trehalose (Figure 1-figure supplement 1C).

**Figure 1:**
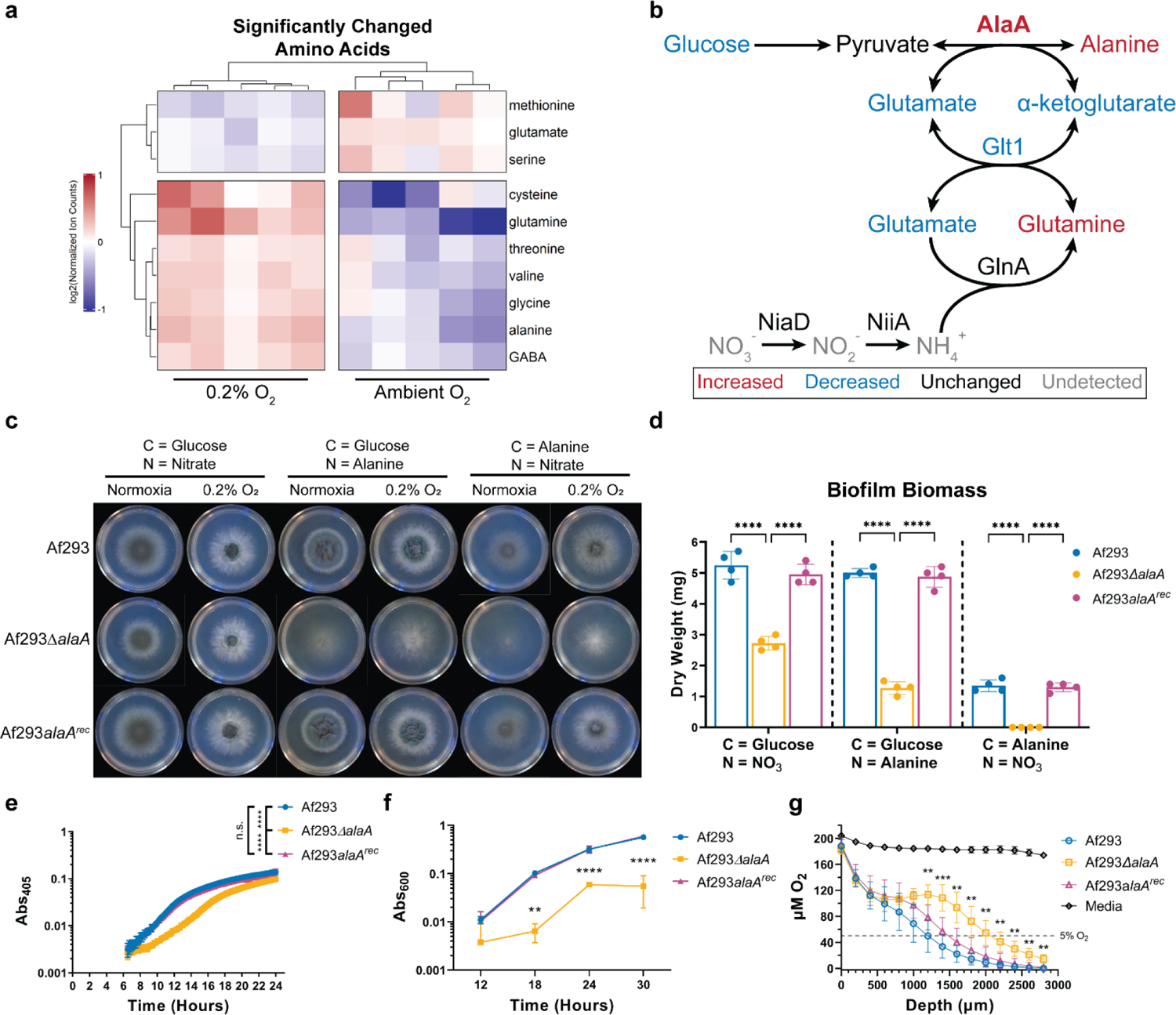
An alanine aminotransferase is required for alanine catabolism and normal biofilm physiology. A) Significantly changed amino acids upon acute exposure to a 0.2% oxygen environment. Ion counts were normalized to the mean ion count for each metabolite across all samples and log_2_ transformed. Each column is a replicate (n = 5 per condition). B) Reaction catalyzed by AlaA and its position in central carbon and nitrogen metabolism. Each metabolite and gene are color-coded according to their relative abundance upon exposure to a 0.2% oxygen environment. Metabolite data was obtained from the experiment in (A), and RNA-sequencing data was obtained from Chung, et al. 2014. C) Growth of Af293Δ*alaA* on minimal media containing the indicated sole carbon and nitrogen sources in ambient oxygen (normoxia) and 0.2% oxygen environments. Images are representative of four replicate cultures. D) Dry biomass of biofilms grown in minimal media containing the indicated sole carbon and nitrogen sources for 24 hours (n = 4). Each replicate is shown along with the mean +/- SD. E) Representative static growth assay of Af293Δ*alaA* over 24 hours of biofilm growth (n = 6 technical replicates). Experiment was repeated at least three times with similar results. F) Crystal violet adherence assay of biofilms grown for 12, 18, 24, and 30 hours (n = 3). G) Oxygen concentration as a function of distance from the air-liquid interface in 24-hour biofilms (n ≥ 7). Culture volumes are approximately 3000µm in depth. ** p < 0.01, *** p < 0.001, **** p < 0.0001, n.s. = not significant by either Two-Way ANOVA with a Tukey’s multiple comparison test (D, F, and G) or One-Way ANOVA with a Tukey’s multiple comparison test (E). All graphs show the mean +/- SD unless otherwise stated.

Intriguingly, many metabolites related to nitrogen metabolism were increased. Specifically, alanine, GABA, glutamine, and several urea cycle intermediates (Figure 1A, Figure 1-figure supplement 1D). This accumulation of nitrogen metabolites could be indicative of either nitrogen storage in the form of more favorable nitrogen sources, as is seen with carbon metabolism described above, and/or potentially indicative of nitrate fermentation as a strategy to recycle reducing potentials and allow glycolysis to continue. The accumulation of alanine was of particular interest due to an association of alanine and low oxygen adaptation in a wide-range of organisms including plants (Lothier et al., 2020; Rocha et al., 2010), crustaceans (Harrison, 2015), flies (Feala et al., 2007), and mammals (Felig et al., 1970). Additionally, several transcriptomic studies involving *A. fumigatus* and oxygen limitation have found that an alanine aminotransferase (Afu6g07770/AFUB_073730) is highly increased in mRNA abundance upon exposure to low oxygen conditions (Figure 1B) (Barker et al., 2012; Chung et al., 2014; Hillmann et al., 2014; Losada et al., 2014). Thus, we further investigated the role of this alanine aminotransferase, herein named *alaA,* in *A. fumigatus* physiology.

### The alanine aminotransferase, *alaA*, is required for efficient catabolism of L-alanine

To assess the role of *alaA* in *A. fumigatus* metabolism and stress resistance, strains lacking the *alaA* gene were generated in the Af293 (Af293Δ*alaA*) and CEA10 (CEA10Δ*alaA*) backgrounds along with respective reconstituted strains in which *alaA* was ectopically re-introduced into the Af293Δ*alaA* and CEA10Δ*alaA* genomes under control of its native promoter (Af293*alaA^rec^* and CEA10*alaA^rec^*). Both Af293Δ*alaA* and CEA10Δ*alaA* colony biofilms grew on glucose minimal media (GMM), where glucose is the sole carbon source and nitrate is the sole nitrogen source, indicating that sufficient alanine was generated for colony biofilm growth independent of *alaA* under these *in vitro* conditions (Figure 1C, Figure 1-figure supplement 2A). However, radial growth of the *alaA* null strains on solid GMM was approximately 10% less than that of their respective WT and reconstituted strains at both ambient O_2_ and 0.2% O_2_ indicating a role for this protein in fungal metabolism in the presence of its preferred carbon source when growing as a colony biofilm (Figure 1C, Figure 1-figure supplement 2A). When the *alaA* null strains were grown with L-alanine as the sole carbon or sole nitrogen source the strains displayed severe colony biofilm growth defects (Figure 1C, Figure 1-figure supplement 2A). Surprisingly, both the wildtype (WT) and the *alaA* null strains grew more robustly at 0.2% O_2_ than ambient O_2_ when alanine was the sole carbon or nitrogen source, despite alanine being a non-fermentable carbon source (Figure 1C). Thus, *alaA* plays an important role in *A. fumigatus* metabolism in multiple carbon, nitrogen, and oxygen environments.

### Robust adherence and growth of *A. fumigatus* biofilms is dependent on *alaA*

*A. fumigatus* submerged and colony biofilms naturally become increasingly oxygen deprived as they mature, albeit to different degrees where submerged biofilms form steeper oxygen gradients than colony biofilms (Kowalski et al., 2020, 2021). We next investigated the role of *alaA* in *A. fumigatus* submerged biofilms. To assess biofilm formation on GMM, and further quantify the role of AlaA in alanine metabolism, we quantified the dry biomass of submerged biofilms grown for 24 hours in GMM and with alanine as a sole carbon or nitrogen source. Loss of *alaA* resulted in a 40-50% decrease in submerged biofilm biomass in GMM. This growth defect was exacerbated when alanine was the sole carbon or nitrogen source, with no biomass recovered when alanine was the sole nitrogen source (Figure 1D). Additionally, we utilized a static growth assay to assess biofilm growth kinetics, which revealed that a*laA* null strains had a longer lag phase than their respective WT or reconstituted strains, indicative of a delay in conidial germination (Figure 1E, Figure 1-figure supplement 2B). To further determine if *alaA* had broad physiological impacts on *A. fumigatus* submerged biofilm formation, a crystal violet adherence assay was utilized to quantify adherence of the *alaA* null biofilms to abiotic surfaces. To account for any impacts of the germination delay on biofilm formation, the adherence of Af293Δ*alaA* was measured over a time course from an immature biofilm at 12 hours to a highly mature biofilm at 30 hours. At all timepoints after 12 hours Af293Δ*alaA* had a severe defect in adherence compared to the WT and reconstituted strains (Figure 1F). CEA10Δ*alaA* was also tested for adherence and showed a similar inability to strongly adhere to surfaces (Figure 1-figure supplement 2C). Finally, we quantified oxygen levels within 24-hour biofilm cultures of Af293Δ*alaA*. The *alaA* null strain cultures were significantly more oxygenated than the WT and reconstituted strains’ biofilms (Figure 1G). However, the portion of the culture containing the bulk of the biofilm’s biomass, depth ∼2000μm - 3000μm based on previous microscopy studies (Kowalski et al., 2020), was still below 5% O_2_ and thereby experiencing hypoxia. Therefore, while the loss of *alaA* has an impact on colony biofilm growth, *alaA* appears to play a greater role in *A. fumigatus* submerged biofilm physiology where steep oxygen gradients naturally occur (Kowalski et al., 2020).

### Catalytic activity of AlaA is required for adherence and alanine growth, but not mitochondrial localization

To begin determining how a gene involved in alanine metabolism is able to impact fungal metabolism and adherence, we generated a catalytically inactive allele of *alaA* to differentiate if the mechanism is through a moonlighting function of the protein or if it is through the catalyzed metabolic reaction. To generate a catalytically inactive allele, the conserved catalytic lysine residue at position 322 (Figure 2-figure supplement 1) (Peña-Soler et al., 2014) was changed to an alanine and a C-terminal GFP tag was added. This construct was then transformed into the native locus of *alaA* in the Af293 background (Af293*alaA^K322A^-GFP*). Additionally, the WT allele was modified with a C-terminal GFP tag and was transformed in the same manner (Af293*alaA-GFP*). Af293*alaA-GFP* and Af293*alaA^K322A^-GFP* grew on alanine as the sole carbon or sole nitrogen source in a manner similar to the WT and *alaA* null strains respectively, confirming that catalytic function was abolished and that the GFP tag did not interfere with protein function (Figure 2A, 2B). A crystal violet adherence assay revealed that Af293*alaA^K322A^-GFP* exhibited an adherence defect equivalent to the deletion of the entire *alaA* gene (Figure 2C). Confocal microscopy of these two strains in combination with Mitotracker^TM^ Deep Red FM revealed that both the WT and catalytically inactive *alaA* alleles were stably expressed and localize to the mitochondria (Figure 2D, Figure 2-figure supplement 2). Mitochondrial localization was surprising given that AlaA lacks a canonical mitochondrial localization signal and may suggest a role in mitochondrial function, as was found to be the case in tumor cells (Beuster et al., 2011). Therefore, AlaA catalytic activity is required for adherence and alanine catabolism, but not mitochondrial localization.

**Figure 2:**
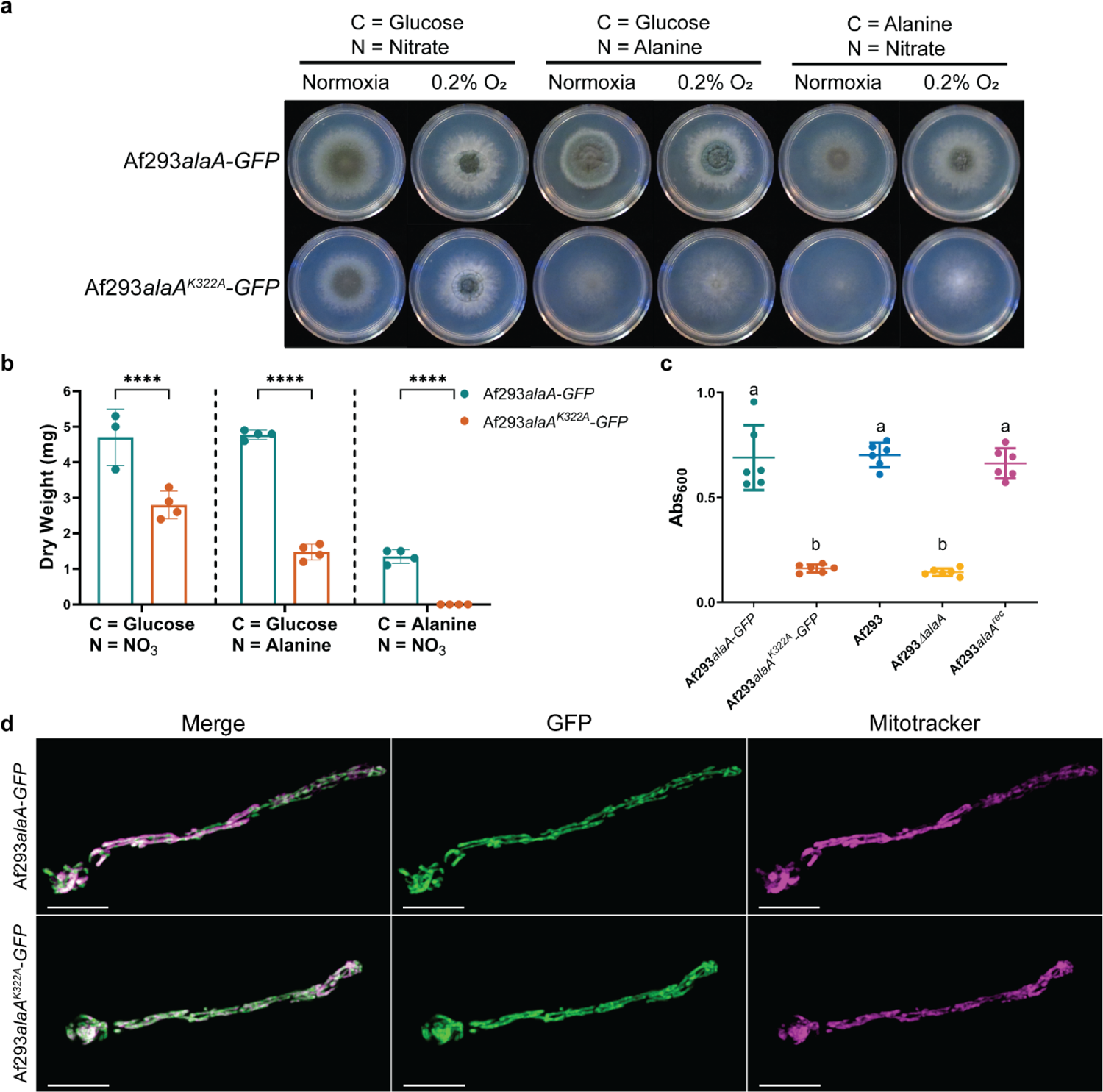
Catalytic activity of AlaA is required for alanine catabolism and adherence of biofilms. A) Growth of Af293*alaA^K322A^-GFP* on minimal media containing the indicated sole carbon and nitrogen sources in ambient oxygen (normoxia) and 0.2% oxygen environments. Images are representative of four replicate cultures. B) Dry biomass of biofilms grown in minimal media containing the indicated sole carbon and nitrogen sources (n ≥ 3). Each replicate along with the mean +/- SD are shown. **** p < 0.0001 as determined by Two-Way ANOVA with a Tukey’s multiple comparisons test. C) Crystal violet adherence assay of 24-hour biofilms (n = 6). Each replicate along with the mean +/- SD are shown. a vs b p < 0.0001 in all comparisons as determined by Two-Way ANOVA with a Tukey’s multiple comparisons test. D) Representative micrographs of germlings containing C-terminal GFP tagged AlaA alleles (green) stained with Mitotracker^TM^ Deep Red FM (magenta).

### Loss of *alaA* leads to alterations in the adherence mediating polysaccharide galactosaminogalactan (GAG)

The primary adherence factor for *A. fumigatus* submerged biofilms studied to date is the extracellular matrix polysaccharide galactosaminogalactan (GAG) (Gravelat et al., 2013; Lee et al., 2015, 2016), and the lack of adherence observed in the *alaA* null strain suggests a role in GAG production or maturation. To examine GAG production, we first utilized a fluorescently labeled lectin specific to N-acetyl-D-galactosamine (GalNAc) residues found in the GAG polysaccharide, FITC-Soybean Agglutinin (SBA). Biofilms of Af293, Af293Δ*alaA*, and Af293*alaA^rec^* were stained with SBA to visualize GAG and calcofluor white, which binds chitin, to visualize biomass at 12, 18, 24, and 30 hours of growth (Figure 3A, D, G). Spinning-disk confocal microscopy was utilized to image the first 300μm of the biofilms, followed by quantification using BiofilmQ (Hartmann et al., 2021). As seen in growth curve experiments (Figure 1F), Af293Δ*alaA* had a lower biovolume at 12 and 18 hours (Figure 3A-C). Total SBA staining of the GAG polysaccharide was quantified as the sum intensity of the SBA stain in each image, revealing that Af293Δ*alaA* biofilms had less total SBA staining than the WT and reconstituted strains starting at 18 hours of growth (Figure 3D-F). While the SBA staining tightly associated with the cell wall at all timepoints in Af293Δ*alaA*, at 30 hours in the WT and reconstituted strains the SBA staining pattern shifted from hyphal associated to primarily staining the extracellular milieu (Figure 3D, G). We quantified the hyphal associated SBA staining as the sum intensity of SBA stain that overlapped with the segmented calcofluor white stain, therefore showing GAG in relation only to hyphal biovolume. In the WT and reconstituted strain biofilms hyphal associated SBA peaked at 18 hours and decreased at 24 and 30 hours as matrix was shed from the hyphae into the extracellular milieu (Figure 3G-I). This was in contrast to total SBA staining, which remained relatively consistent from 18-30 hours of growth in the WT and reconstituted strains (Figure 3E).

**Figure 3:**
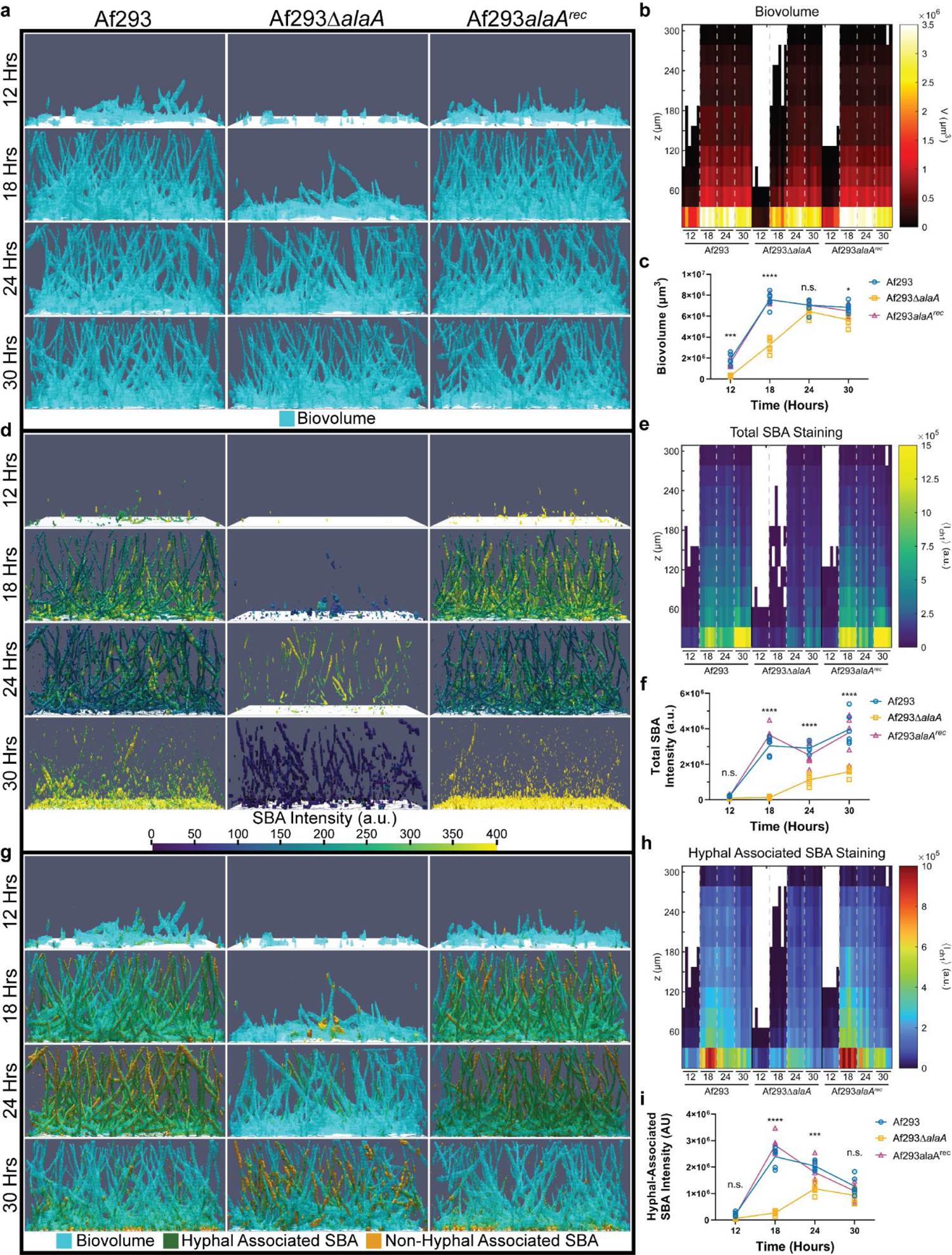
Loss of *alaA* alters extracellular matrix staining by the galactosaminogalactan binding lectin SBA. A) Representative image renderings of biovolume in the first 300µm of biofilms grown for 12, 18, 24, and 30 hours. Biofilms were stained with calcofluor white and FITC-SBA followed by fixing with paraformaldehyde. Biovolume was determined by segmentation of the calcofluor white stain of each image. B) Heatmap of biovolume as a function of height from the base of the biofilm. C) Global segmented biovolume quantifications of each biofilm. D) Representative image renderings of FITC-SBA staining intensity corresponding to biomass images in (A). Renderings show FITC-SBA matrix intensity mapped onto the segmented FITC-SBA stain. E) Heatmap of FITC-SBA intensity as a function of height from the base of the biofilm. F) Sum intensity quantification of FITC-SBA staining for each biofilm. G) Representative merged image renderings of the segmented biovolume (calcofluor white), shown in blue, and segmented FITC-SBA stain shown in orange. Hyphal associated SBA staining will appear green as a result of the overlap between the two channels. SBA was considered hyphal-associated or non-hyphal associated based on overlap with the segmented biomass. H) Heatmap of hyphal associated FITC-SBA intensity as a function of height from the base of the biofilm. I) Sum intensity quantification of hyphal associated FITC-SBA staining for each biofilm. Each graph and heatmap shows the individual replicates for each timepoint (n = 6). For (C, F, and I), the line goes through the mean of each timepoint. * p < 0.05, *** p < 0.001, **** p < 0.0001, n.s. = not significant as determined by Two-Way ANOVA with a Tukey’s multiple comparison’s test for (C, F, and I).

**Figure 4:**
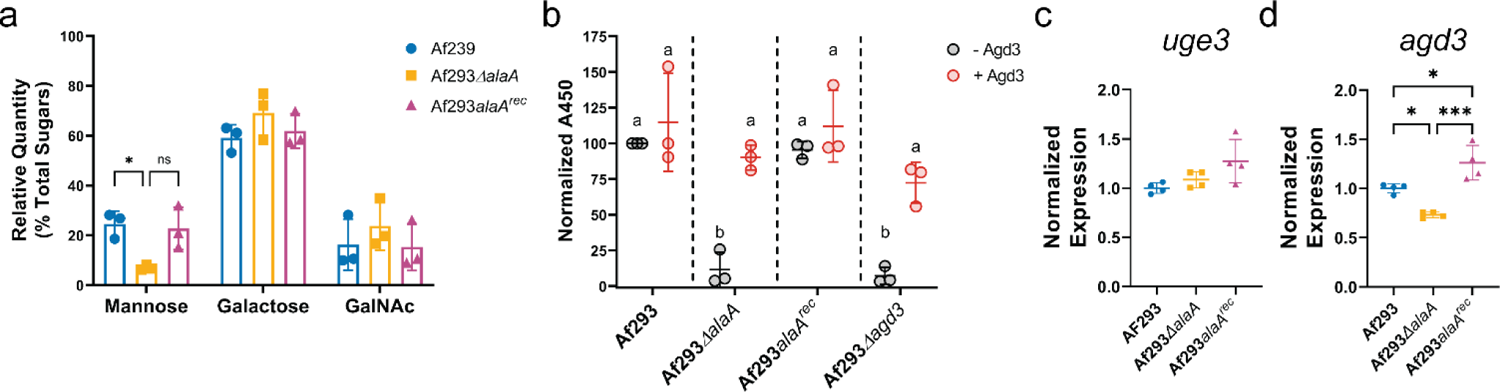
*alaA* is required for proper deacetylation of galactosaminogalactan. A) Monosaccharide analysis of extracellular matrix polysaccharides (n = 3). * p < 0.05, n.s. = not significant by Two-Way ANOVA with a Tukey’s multiple comparisons test. B) Enzyme linked lectin assay (ELLA) of biofilm extracellular matrix treated or untreated with recombinant Agd3. a vs b p < 0.05 for all comparisons as determined by Two-Way ANOVA with a Tukey’s multiple comparisons test. C-D) Expression of *uge3* (C) and *agd3* (D) in 24-hour biofilm cultures as determined by RTqPCR. * p < 0.05, *** p < 0.001 as determined by One-Way ANOVA with a Tukey’s multiple comparisons test for C and D. For all graphs each replicate along with the mean +/- SD is shown.

To chemically define how GAG was being altered in the *alaA* null strain, monosaccharide analysis of ECM polysaccharides and an enzyme-linked lectin assay (ELLA) were conducted. Monosaccharide analysis revealed that the *alaA* null strain’s ECM polysaccharide composition was similar to that of the WT strain (Figure 4A). This finding suggests that the altered ECM is primarily due to a change in the maturation of the ECM polysaccharides rather than a difference in the base polysaccharides produced. After GAG has been synthesized, partial deacetylation by the Agd3 deacetylase is necessary for functional adherence (Bamford et al., 2020; Lee et al., 2016). To test if GAG maturation was altered, we utilized an ELLA in combination with treatment of the ECM by recombinant Agd3. In principle, deacetylated GAG in supernatants allows for adherence to the walls of a polystyrene plate, whereas fully acetylated GAG is unable to adhere and is easily removed by washing. Adherent, and therefore deacetylated, GAG can then be quantified by binding of a biotinylated SBA lectin coupled to a streptavidin-conjugated horseradish peroxidase. Additionally, the presence of fully acetylated GAG can be detected by pre-treating samples with recombinant Agd3, producing de-acetylated, adherent, GAG that can then be detected by SBA. A strain lacking the *agd3* gene, which only produces fully acetylated GAG (Lee et al., 2016), was utilized as a control. The *alaA* and *agd3* null strains both yielded low levels of adherent (deacetylated) GAG compared to the WT, and this was rescued by treatment of ECM with recombinant Agd3 protein (Figure 4B). Therefore, *alaA* is not required for GAG production, but rather is required for deacetylation and maturation of the GAG polysaccharide into its functional form. Finally, mRNA abundance of *uge3* and *agd3* was measured from RNA isolated from 24-hour biofilms of Af293, Af293Δ*alaA*, and Af293*alaA^rec^* to begin to distinguish if the observed differences in GAG are through a transcriptional or post-transcriptional mechanism of regulation. No differences in expression of *uge3* were observed (Figure 4C). While a statistically significant decrease of ∼20% in *agd3* mRNA levels was observed (Figure 4D), it is unclear if that level of mRNA difference could cause the degree of altered GAG deacetylation observed. Thus, while loss of *alaA* has a modest impact on *agd3* at the transcriptional level, the impact of *alaA* on GAG maturation is likely to be primarily post-transcriptional.

### Deletion of *alaA* leads to cell wall changes and increased susceptibility of biofilms to echinocandins

Given that GAG maturation was substantially impacted by loss of *alaA* and that *alaA* plays a role in metabolism, we asked if loss of *alaA* impacts additional cell wall polysaccharides. To test this hypothesis, germlings of Af293, Af293Δ*alaA*, and Af293*alaA^rec^* were stained with calcofluor white (to measure total chitin), wheat-germ agglutinin (WGA) (to measure surface exposed chitin), and soluble Dectin-1 Fc (to measure surface exposed β-glucans). Curiously, Af293Δ*alaA* germlings had lower WGA staining (exposed chitin), despite no difference in calcofluor white staining (total chitin) (Figure 5A-B). Loss of *alaA* also decreased exposure of the immunostimulatory β-glucan polysaccharide, as determined by Dectin-1 Fc staining (Figure 5C). These results were surprising, as it has been shown that GAG masks β-glucans, and perturbations to GAG synthesis (Gravelat et al., 2013) or maturation (Lee et al., 2016) normally result in higher levels of Dectin-1 Fc staining. Together this suggests Af293Δ*alaA* has lower levels of total cell wall β-glucans and that *alaA* is required for WT cell wall organization.

**Figure 5:**
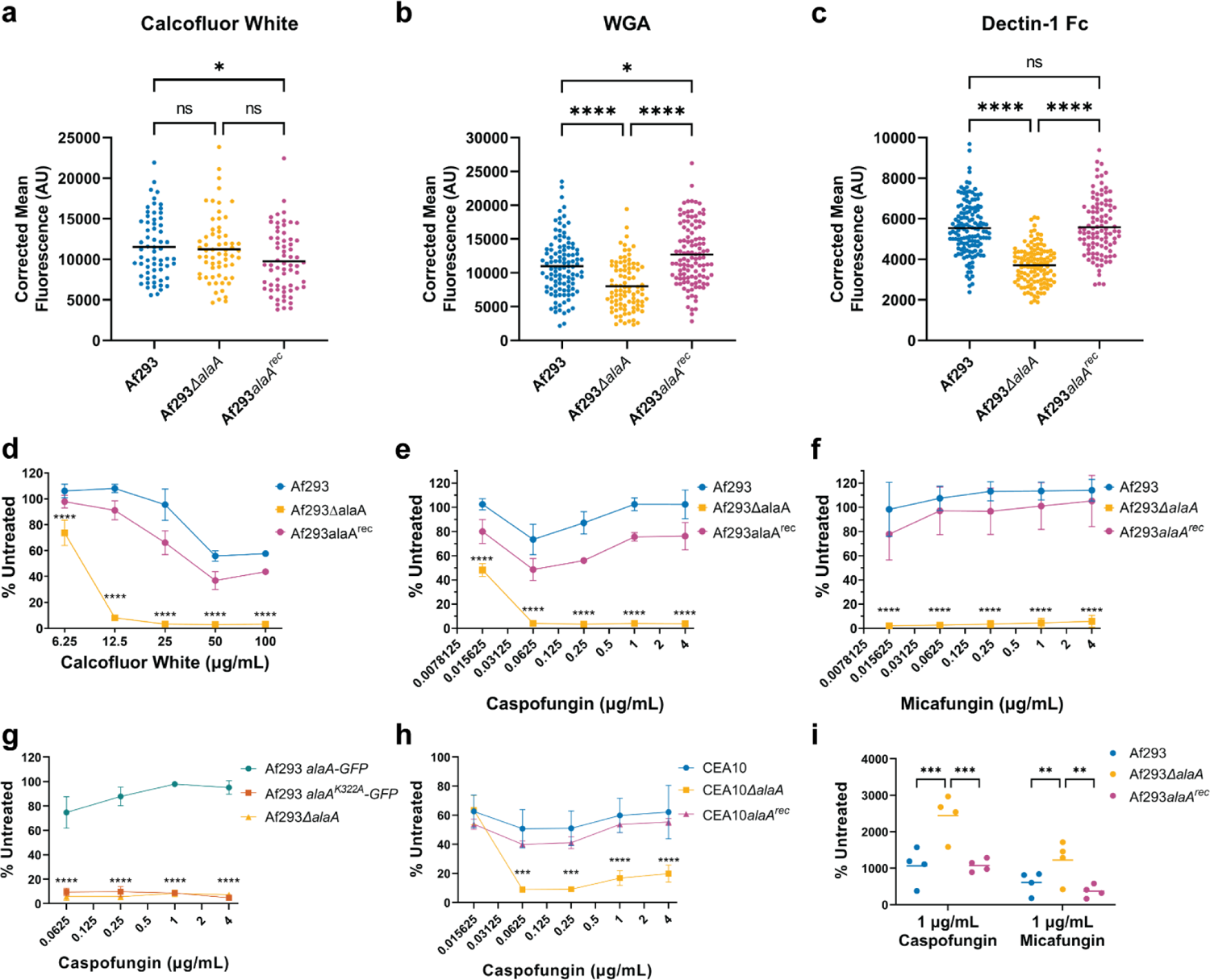
Loss of *alaA* leads to cell wall changes and increased susceptibility of biofilms to echinocandins. Germlings were stained with calcofluor white to quantify total chitin content (A), FITC-wheat germ agglutinin (WGA) to quantify surface exposed chitin (B), and Dectin-1 Fc to quantify surface exposed β-glucans (C). Each data-point represents an individual germling across three independent cultures per strain for each cell wall stain and the lines correspond to the mean. * p < 0.05, **** p < 0.0001, n.s. = not significant as determined by One-Way ANOVA with a Tukey’s multiple comparisons test. D-F) 24-hour biofilms were established in the absence of drug and treated with calcofluor white (D), caspofungin (E), or micafungin (F) at the indicated concentrations for 3 hours and viability was determined by XTT assay. Mean +/- SD are shown for n ≥ 3 for each experiment. **** p < 0.0001 as determined by Two-Way ANOVA with a Tukey’s multiple comparison test. The highest p-value for Af293Δ*alaA* compared to both Af293 and Af293*alaA^rec^* is shown. G) Af293*alaA^K322A^-GFP* biofilms were grown for 24-hours, treated with caspofungin at the indicated concentrations for 3 hours, and viability was determined by XTT assay. Mean +/- SD are shown for n = 3 replicates. **** p < 0.0001 as determined by Two-Way ANOVA with a Tukey’s multiple comparison test. The highest p-values for Af293*alaA^K322A^-GFP* and Af293Δ*alaA* in comparison to Af293*alaA-GFP* are shown. No significant difference was observed between Af293*alaA^K322A^-GFP* and Af293Δ*alaA*. H) CEA10Δ*alaA* biofilms were grown for 24-hours, treated with caspofungin at the indicated concentrations for 3 hours, and viability was determined by XTT assay. Mean +/- SD are shown for n = 3 replicates. *** p < 0.001, **** p < 0.0001 as determined by Two-Way ANOVA with a Tukey’s multiple comparison test. The highest p-values for CEA10Δ*alaA* compared to both CEA10 and CEA10*alaA^rec^* are shown. I) Adenylate kinase release assay as a quantification of cell lysis. 24-hour biofilms were treated with 1µg/mL of caspofungin (left) or micafungin (right) for 3-hours and supernatant adenylate kinase activity was quantified. Each replicate and the mean are shown (n = 4). ** p < 0.01, *** p < 0.001 as determined by Two-Way ANOVA with a Tukey’s post-test.

To determine if these cell wall changes translated to functional phenotypes, biofilms of Af293, Af293Δ*alaA*, and Af293*alaA^rec^* were tested for sensitivity to the cell wall perturbing agents calcofluor white and the echinocandin class of antifungal drugs. Biofilms were grown to maturity (24 hours) prior to application of cell wall stress for 3 hours. Viability was then compared to untreated biofilms via reduction of the metabolic dye XTT. Af293Δ*alaA* biofilms were significantly more susceptible to damage by calcofluor white at all concentrations tested, with greater than 90% inhibition beginning at 12.5 μg/mL (Figure 5D). In contrast, the WT and reconstituted strains maintained at least 30% viability at even the highest concentration tested (100 μg/mL) (Figure 5D). We next tested the strains for susceptibility to the echinocandin caspofungin. The WT and reconstituted biofilms displayed minimal damage regardless of concentration. Quantification also reveals signs of the paradoxical effect in this system, where the fungus will recover growth as the concentration of drug increases beyond a MEC (Figure 5E). Thus, similar to MEC and agar colony biofilm plate assays that begin with conidia, mature WT *A. fumigatus* biofilms are caspofungin tolerant. However, *alaA* null biofilms were highly susceptible to caspofungin and reached >90% inhibition at a concentration of 0.0625 μg/mL. Unlike WT biofilms, the *alaA* null biofilms did not display evidence of a paradoxical effect, with increasing concentrations of caspofungin yielding equivalent or greater damage (Figure 5E).

To test if this increased susceptibility to caspofungin extended to other echinocandins, these experiments were validated with another echinocandin, micafungin. Similar to caspofungin treatment, the biofilms of WT and reconstituted strains were highly resistant to treatment with micafungin, whereas Af293Δ*alaA* was inhibited >90% at even the lowest concentration of drug tested, 0.015625 μg/mL (Figure 5F). The catalytically inactive strain (Af293*alaA^K322A^*) was also tested for caspofungin sensitivity and displayed the same phenotype as Af293Δ*alaA* (Figure 5G). Finally, to ensure this phenotype was not specific to the Af293 reference strain, these phenotypes were validated in another reference background, CEA10. CEA10, CEA10Δ*alaA*, and CEA10*alaA^rec^* biofilms were tested for susceptibility to caspofungin and again the loss of *alaA* resulted in increased echinocandin susceptibility (Figure 5H). Moreover, the increased susceptibility of the *alaA* null mutant was confirmed by measuring adenylate kinase release (Didone et al., 2011). In brief, adenylate kinase is normally present in very low quantities extracellularly and increased release of adenylate kinase is indicative of cell lysis. Treatment of Af293Δ*alaA* biofilms with caspofungin or micafungin resulted in a greater release of adenylate kinase into the supernatant, indicating that loss of *alaA* potentiates the fungicidal effects of echinocandins (Figure 5I, FIGURE 5-FIGURE SUPPLEMENT 1). Thus, the presence and catalytic activity of AlaA are required for the high level of echinocandin resistance observed in phylogenetically diverse *A. fumigatus* biofilms.

We next tested if the increased susceptibility of Af293Δ*alaA* to echinocandins was biofilm specific or if it extended to more traditional measures of drug resistance and tolerance that begin with exposing dormant conidia to the drug. Tolerance, or the ability to grow in the presence of a fixed concentration of drug, to caspofungin was measured using radial growth of conidia on agar plates at three concentrations of caspofungin (0.25 μg/mL, 1 μg/mL, and 4 μg/mL). No significant differences in colony biofilm growth were observed between the strains at any concentration tested on agar surfaces (Figure 5-figure supplement 2A-C). Additionally, the paradoxical effect was observed in all three strains tested, with increased growth as the concentration of caspofungin increased. Intriguingly, no difference in resistance to caspofungin was observed when a minimal effective concentration (MEC) assay was utilized with conidia of Af293, Af293Δ*alaA*, and Af293*alaA^rec^* (Figure 5-figure supplement 2D). Therefore, the increased susceptibility of Af293Δ*alaA* to echinocandins is a biofilm specific phenomenon, as no difference in resistance or tolerance to caspofungin is observed when the drug is applied to dormant conidia.

### Chemical Inhibition of AlaA by β-chloro-L-alanine decreases adherence and increases susceptibility of *A. fumigatus* biofilms to caspofungin

Given the potential clinical significance of increasing *A. fumigatus* biofilm susceptibility to echinocandins, we next tested whether chemical inhibition of AlaA was sufficient to confer similar phenotypes observed in the null or catalytically inactive mutant strains. The chemical β-chloro-L-alanine has previously been shown to inhibit mammalian alanine aminotransferases (Beuster et al., 2011; Golichowski & Jenkins, 1978; Morino et al., 1979), and thus we tested if β-chloro-L-alanine treatment could recapitulate the adherence and caspofungin phenotypes observed in the *alaA* null strain. Af293, Af293Δ*alaA*, and Af293*alaA^rec^* were incubated with 10-fold increasing concentrations of β-chloro-L-alanine from 0.1μM to 1000μM and tested for adherence via crystal violet adherence assay. Increasing concentrations of β-chloro-L-alanine resulted in decreased adherence for Af293 and Af293*alaA^rec^*, with EC_50_ values of 10.49 μM and 15.90 μM respectively (Figure 6A). At 100μM of β-chloro-L-alanine, adherence of the WT and reconstituted strains was inhibited to slightly above that of the *alaA* null strain. Importantly, adherence of the Af293Δ*alaA* was unaltered by any concentration of β-chloro-L-alanine tested indicating some level of chemical specificity for AlaA (Figure 6A). Additionally, treatment of the GFP-tagged AlaA and catalytically inactive strains with β-chloro-L-alanine yielded similar results to the WT and *alaA* null strains, respectively, indicating that the compound is acting through the catalytic activity of the AlaA enzyme (Figure 6B).

**Figure 6:**
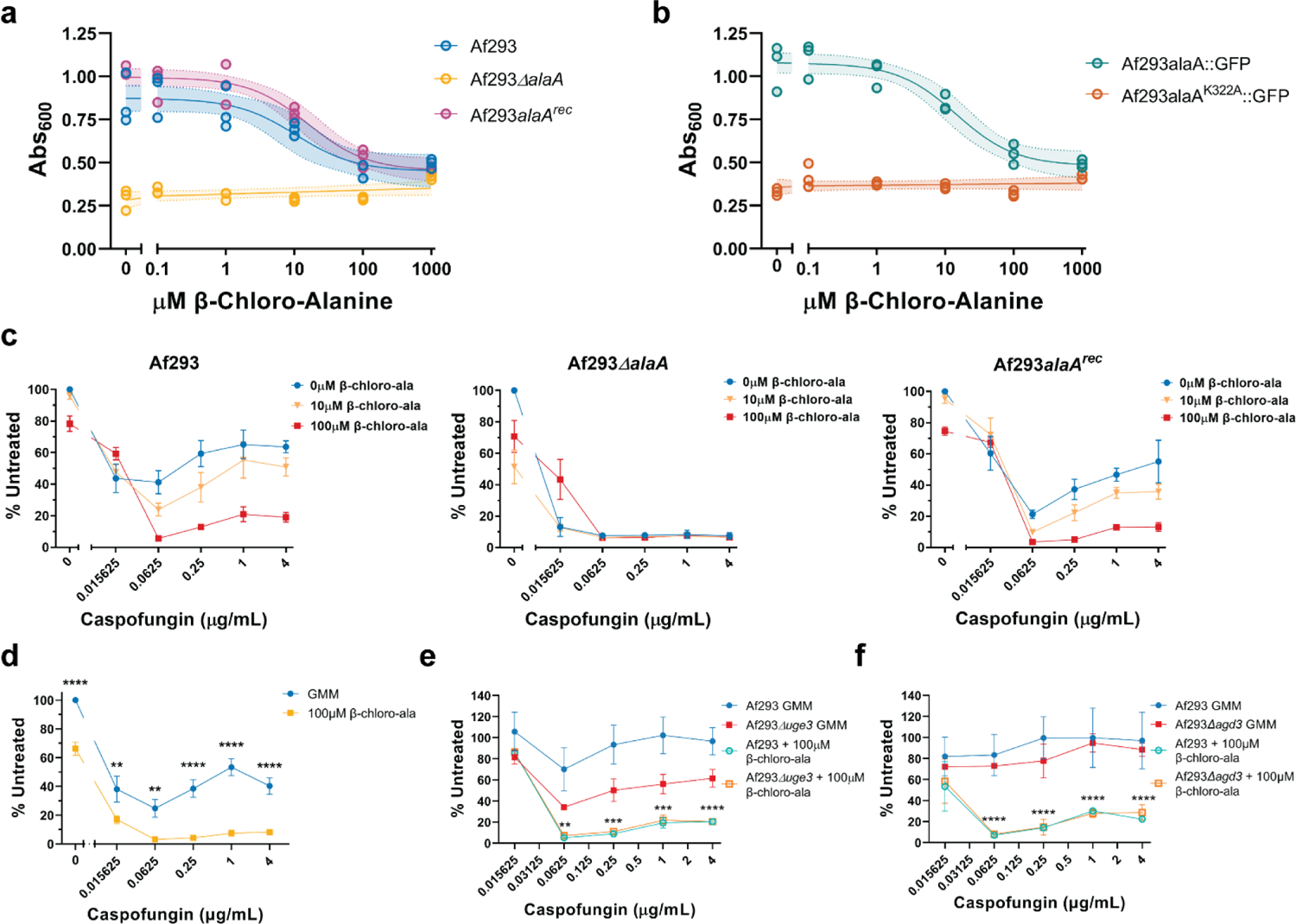
Chemical inhibition of AlaA by β-chloro-L-alanine is sufficient to decrease adherence and increase susceptibility of biofilms to caspofungin. A-B) Crystal violet adherence assay of 24-hour biofilms grown in the presence of increasing concentrations of β-chloro-L-alanine. Each replicate is shown (n = 3) along with a non-linear regression using a dose-response model (line) +/- 95% confidence interval (shaded area). C) Susceptibility of 24-hour biofilms of Af293 (left), Af293Δ*alaA* (middle), and Af293*alaA^rec^* (right) established in the presence of 0, 10, or 100µM β-chloro-L-alanine. Biofilms were treated with the indicated concentrations of caspofungin for 3-hours and viability was assessed by XTT assay. Mean +/- SD are shown (n = 3). D) Susceptibility of 24-hour CEA10 biofilms established in the presence or absence of 100µM β-chloro-L-alanine to caspofungin treatment. Biofilms were treated with the indicated concentrations of caspofungin for 3-hours and viability was assessed by XTT assay. Mean +/- SD are shown (n = 3). ** p < 0.01, **** p < 0.0001 as determined by Two-Way ANOVA with Tukey’s multiple comparisons test. E-F) Susceptibility of 24-hour Af293Δ*uge3* (E) and Af293Δ*agd3* (F) biofilms established in the presence or absence of 100µM β-chloro-L-alanine to caspofungin treatment. Biofilms were treated with the indicated concentrations of caspofungin for 3-hours and viability was assessed by XTT assay. Mean +/- SD are shown (n = 3). ** p < 0.01, *** p < 0.001, **** p < 0.0001 as determined by Two-Way ANOVA with a Tukey’s multiple comparisons test. The highest p-values for β-chloro-L-alanine treated groups vs their respective untreated groups are shown.

To test if β-chloro-L-alanine treatment increases susceptibility of biofilms to echinocandins, biofilms were grown in the presence of 10μM and 100μM β-chloro-L-alanine to represent a range of values that encompass both the EC_50_ value determined by the adherence assay results (10μM) and the concentration that yielded an *alaA* deletion-like phenotype (100μM) (Figure 6A). The β-chloro-L-alanine treated biofilms of Af293, Af293Δ*alaA*, and Af293*alaA^rec^* were tested for sensitivity to caspofungin treatment. In the WT and reconstituted strains efficacy of caspofungin increased as the concentration of β-chloro-L-alanine increased (Figure 6C). Curiously, even in the *alaA* null strain basal XTT reduction decreased as the concentration of β-chloro-L-alanine increased. However, the *alaA* null strain remained highly susceptible to caspofungin regardless of the concentration of β-chloro-L-alanine (Figure 6C). Additionally, treatment of CEA10 with β-chloro-L-alanine increased caspofungin susceptibility of biofilms validating that this is not specific to the Af293 strain background (Figure 6D). Together these data establish the proof of concept that chemical inhibition of AlaA is a possible strategy for increasing susceptibility of *A. fumigatus* biofilms to echinocandins.

### Altered extracellular matrix is not the primary factor impacting caspofungin susceptibility

In other fungi, the biofilm extracellular matrix has been shown to be a major factor in reducing antibiotic efficacy against biofilms (Taff et al., 2013). Therefore, we wanted to test if the increased susceptibility to caspofungin could be attributed to the altered GAG composition of the *alaA* null strain. To do this we utilized a strain lacking the UDP-glucose-4-epimerase required to produce GAG (Af293Δ*uge3*) and a strain lacking the deacetylase required for the maturation of GAG (Af293Δ*agd3*) in combination with β-chloro-L-alanine treatment (Gravelat et al., 2013; Lee et al., 2016). In an attempt to overcome the inability of Af293Δ*uge3* and Af293Δ*agd3* to adhere to abiotic surfaces (Gravelat et al., 2013; Lee et al., 2016), we performed experiments with these strains on collagen-coated tissue culture plates. Collagen is a mammalian extracellular matrix component abundant in the lung, and we observed that this treatment was sufficient to partially restore adherence of both strains, indicating the existence of GAG-independent mechanisms of adherence to alternative substrates found in mammalian lungs (Figure 6-figure supplement 1). Af293, Af293Δ*uge3*, and Af293Δ*agd3* biofilms were established on collagen-coated plates with or without 100μM β-chloro-L-alanine and then subsequently treated with caspofungin. All three strains were highly susceptible to caspofungin when AlaA was inhibited by β-chloro-L-alanine (Figure 6E, 6F). Untreated Af293Δ*uge3* was inhibited by caspofungin treatment to a greater extent than the WT strain. However, this increased susceptibility was far less severe than observed in β-chloro-L-alanine treated biofilms and was not observed in the deacetylase deficient strain (Af293Δ*agd3*) (Figure 6E). Therefore, GAG contributes to caspofungin resistance to some degree, but it is not the primary factor responsible for the increased susceptibility when AlaA is chemically inhibited or genetically altered.

### *alaA* is required for echinocandin resistance *in vivo*

Finally, we sought to determine if *alaA* plays a role in echinocandin resistance *in vivo* within lung infection microenvironments. To address this question, we utilized a chemotherapy murine model of invasive pulmonary aspergillosis (IPA). Outbred CD1 mice were immunosuppressed with cyclophosphamide and triamcinolone then challenged with conidia of Af293, Af293Δ*alaA*, or Af293*alaA^rec^*. The infection was allowed to establish for 24 hours followed by three treatments with either 0.9% NaCl or 1mg/kg micafungin every 24 hours (Figure 7A). 12 hours after the final micafungin treatment relative fungal burden was determined by qPCR quantification of *A. fumigatus* 18S rDNA. The dosage of micafungin treatment used had no significant impact on fungal burden in mice inoculated with the WT or reconstituted strains. Moreover, loss of *alaA* at the time point examined did not significantly impact fungal burden levels in the untreated groups (Figure 7B). In contrast, there was a 4-fold reduction in fungal burden in mice inoculated with Af293Δ*alaA* and treated with micafungin compared to untreated mice (Figure 7B). Thus, loss of *alaA in vivo* significantly increases the susceptibility of *A. fumigatus* to sub-effective concentrations of the echinocandin micafungin.

**Figure 7:**
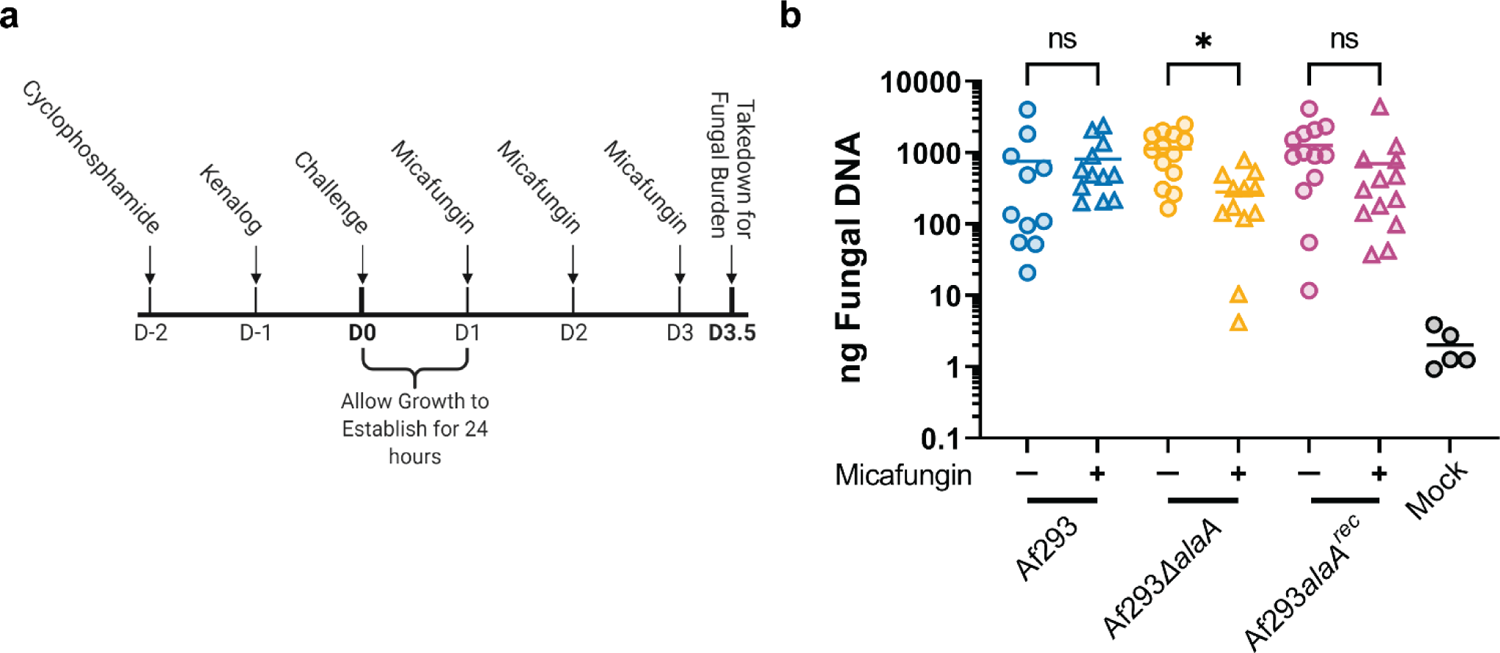
*alaA* is required for echinocandin resistance *in vivo*. A) Experimental outline for determining *in vivo* echinocandin resistance using a chemotherapy model of invasive aspergillosis. Outbred CD1 mice were immunosuppressed with 150mg/kg cyclophosphamide 48 prior to fungal challenge and 40mg/kg triamcinolone 24 hours prior to fungal challenge. Mice were challenged with *A. fumigatus* or PBS (mock) at D0, and infection was allowed to establish for 24 hours. Mice were treated with either 1mg/kg micafungin or 0.9% NaCl every 24 hours from D1 to D3. 12 hours after the final micafungin treatment mice were sacrificed for fungal burden determination by qPCR. B) qPCR quantification of total ng fungal DNA in lungs of mice challenged with the indicated *A. fumigatus* strains and treated or untreated with 1mg/kg micafungin according to design in (A). Each datapoint and the mean are shown (n ≥ 12 for each experimental group and n = 5 for mock infected mice across two independent experiments). * p < 0.05, n.s. = not significant as determined by Kruskal-Wallis with a Dunn’s multiple comparisons test.

## Discussion

The vast majority of antifungal drug studies and discoveries are based on treatment of the conidial state of *A. fumigatus* despite the fact that clinical treatment is largely in the context of an already established infection where *A. fumigatus* exists as a biofilm. Here we discovered a gene involved in the metabolism of alanine that confers biofilm specific echinocandin resistance to *A. fumigatus*. Given the results presented in this work, as well as others, we have shown that antifungal treatment of biofilms is far less effective than treatment of conidia (Kowalski et al., 2020; Seidler et al., 2008). Mechanisms to increase efficacy of antifungal drugs in the context of *A. fumigatus* biofilms are of great clinical need and may even bridge the discrepancies between relatively low, but rising, rates of antifungal resistance *in vitro* and the relatively high rates of clinical treatment failure (Herbrecht et al., 2015; Maertens et al., 2016, 2021; Marr et al., 2015). Both genetic and chemical disruption of AlaA results in increased susceptibility of biofilms to echinocandin treatment, suggesting inhibition of this enzyme is a potential treatment option for clinically enhancing the efficacy of this class of antifungals. While alanine aminotransferase inhibitors have already been developed (Beuster et al., 2011; Golichowski & Jenkins, 1978; Morino et al., 1979), these molecules were discovered using mammalian alanine aminotransferases (AlaA has 45.77% and 45.19% amino acid sequence identity with human GPT1 and GTP2, respectively (Figure 2-figure supplement 1B-C)). Thus, the chemical inhibition performed in this study serves as a proof of principle for targeting AlaA activity in a combination therapy approach. However, these molecules could provide chemical building blocks for development of more fungal specific and/or potent alanine aminotransferase inhibitors. β-chloro-L-alanine has been utilized in a murine cancer model where inhibition of murine alanine aminotransferase was found to decrease the Warburg effect and increase mitochondrial activity of tumor cells (Beuster et al., 2011). While this is promising for future studies *in vivo*, further safety and pharmacological studies are needed.

We originally found *alaA* through investigation of datasets associated with low oxygen adaptation. AlaA catalyzes interconversion of pyruvate and alanine without direct involvement of reducing potentials or any high energy molecules, such as ATP. However, the reduction of nitrate to alanine would consume five reducing potentials and this pathway is suggested to be important for a variety of systems in the adaptation to low oxygen (Feala et al., 2007; Felig et al., 1970; Harrison, 2015; Lothier et al., 2020; Rocha et al., 2010). It is possible that in these systems alanine serves as a nitrogen sink to prevent toxic ammonium accumulation during the conversion of nitrate to ammonium. Therefore, we had originally hypothesized that AlaA function was a critical means of recycling reducing potentials during low oxygen growth and were surprised to find that AlaA plays a significant role in polysaccharide regulation and biofilm formation despite the minimal impact on growth in a low oxygen environment. This result could be due to a high redundancy in the number of mechanisms encoded by the fungus to balance reducing potentials, or it could suggest that alanine metabolism has a more specific role in adaptation to natural oxygen gradients formed by respiration and/or adaptation to stochastic fluctuations in environmental oxygen that naturally occur during filamentous fungal biofilm growth.

The involvement of an alanine aminotransferase in polysaccharide regulation is even more perplexing when one considers that none of the components of the reaction (alanine, pyruvate, glutamate, and α-ketoglutarate) have any obvious or direct role in any known biochemical pathways to generate *A. fumigatus* cell wall or extracellular matrix components. AlaA also does not appear to be essential for maintaining basal levels of alanine for protein production, as genetic disruption of *alaA* did not yield alanine auxotrophy. Here we describe that the catalytic activity of the AlaA protein is essential for polysaccharide regulation. Additionally, transcriptional data corresponding to the *uge3* and *agd3* genes essential for synthesis and maturation of GAG, respectively, suggests that the mechanism of regulation is predominantly post-transcriptional and potentially indirect from alterations in fungal metabolism (Figure 4C-D). However, the exact mechanism through which this reaction regulates polysaccharide biosynthesis and maturation remains to be further studied. Further investigation into this mechanism could yield significant insight into the interplay between metabolism, biofilm formation, and antifungal drug resistance to help inform development of novel biofilm targeted antifungal drugs.

## Materials and Methods

### Strains and growth conditions

Mutant strains were made in the *A. fumigatus* Af293 and CEA10 strains, and therefore Af293 and CEA10 were used as the wildtype (WT) strains as appropriate for each experiment. Strains were stored as conidia in 25% glycerol at −80°C and maintained on 1% glucose minimal media (GMM; 6g/L NaNO3, 0.52g/L KCL, 0.52g/L MgSO4•7H2O, 1.52g/L KH2PO4 monobasic, 2.2mg/L ZnSO4•7H20, 1.1mg/L H3BO3, 0.5mg/L MnCl2•4H2O, 0.5mg/L FeSO4•7H2O, 0.16mg/L CoCl2•5H2O, 0.16mg/L CuSO4•5H2O, 0.11mg/L (NH4)6Mo7O24•4H2O, 5mg/L Na4EDTA, 1% glucose; pH 6.5). Solid media was prepared by addition of 1.5% agar. All experiments were performed with GMM unless explicitly stated otherwise. For experiments where alanine was the sole carbon or nitrogen source, glucose or NaNO_3_ were replaced with alanine at an equimolar quantity of carbon or nitrogen atoms, respectively. For all experiments, *A. fumigatus* was grown on solid GMM at 37°C 5% CO_2_ for 3 days to produce conidia. Conidia were collected using 0.01% Tween-80, counted using a hemacytometer, and diluted in either 0.01% Tween-80 or media to the final concentration used in each assay.

### Strain construction

*alaA* null mutants were generated by replacing the *alaA* open-reading frame (AFUB_073730/Afu6g07770) with the dominant selection marker *ptrA* in both the Af293 and CEA10 backgrounds. The replacement construct was generated using overlap PCR to fuse ∼1kb upstream and ∼1kb downstream of the open reading frame of *alaA* to the *ptrA* marker. The resulting construct was transformed into protoplasts of each strain and mutants were selected for on osmotically stabilized minimal media (GMM plus 1.2M sorbitol) containing 100 μg/L pyrithiamine hydrobromide (Sigma). Reconstitution of the *alaA* gene was performed by PCR amplification of the *alaA* locus for each strain from ∼1.1kb upstream of the start codon to ∼700bp downstream of the stop codon using primers containing PacI and AscI digestion sites. The resulting PCR products were digested with PacI and AscI restriction enzymes and individually ligated into a plasmid containing the hygromycin resistance marker

*hygR*. The resulting plasmids were ectopically transformed into protoplasts derived from the *alaA* null strain for each plasmid’s respective background. Reconstituted mutants were selected for on osmotically stabilized minimal media containing both 175μg/mL hygromycin B (VWR) and 100μg/L pyrithiamine to ensure the mutated locus remained intact. GFP tagged alleles of WT *alaA* and catalytically inactive *alaA^K322A^* were generated at the *alaA* native locus in the Af293 background using the *ptrA* marker. The WT allele was generated using overlap PCR to fuse ∼1kb upstream of the stop codon of alaA, excluding the stop codon, to a fragment containing an in-frame *gfp* linked to a *trpC* terminator from *A. nidulans* and the *ptrA* marker, along with the same ∼1kb downstream of the stop codon that was used in the deletion construct. The catalytically inactive mutation was generated using nested PCR from the mutation site to immediately before the stop codon in order to modify the AAG lysine codon to a GCC alanine codon. This fragment was then fused with 500bp upstream of the point mutation, along with the in frame gfp-trpC_terminator_ *ptrA* fragment, and ∼1kb downstream of the *alaA* stop codon. The two alleles were transformed into Af293 protoplasts and mutants were selected using pyrithiamine. Sanger sequencing was used to confirm each allele.

Protoplasts were generated using lysing enzyme from *Trichoderma harzianum* (Sigma) and transformed as previously described (Willger et al., 2008). Protoplasts were plated on sorbitol stabilized minimal media (GMM + 1.2M sorbitol) containing pyrithiamine. For hygromycin selection protoplasts were allowed to recover without hygromycin selection until germtubes were visible by inverted microscope (overnight at 37°C). At which point 0.6% agar media containing hygromycin was added to a final concentration of 175μg/mL. All strains were single spored and checked for correct integration, or presence of construct in the case of the ectopic reconstituted strains, via PCR and southern blot. Additionally, the basal expression of *alaA* was checked by RT-qPCR on RNA extracted from 24-hour biofilms for the reconstituted strains using the *alaA* null mutants as negative controls (Figure 1-figure supplement 3).

### Metabolomics

Cultures for metabolomics were performed with 100mL of 10^6 conidia/mL of CEA10. Shaking liquid cultures were performed in baffled flasks with a foam stopper to allow rapid environmental acclimation to changes in oxygen tension. Cultures were grown for 24 hours at 37°C 200RPM in ambient oxygen followed by either continued incubation at ambient oxygen or a shift to 0.2% O_2_ for two hours. Biomass was harvested by filtering through Miracloth, washed thoroughly with water, and flash frozen in liquid nitrogen. The biomass was then lyophilized and 100mg dry weight was submitted to Metabolon for LC-MS/MS analysis, metabolite identification, and relative quantification. Data processing and figure generation was performed in R using the ComplexHeatmap (Gu et al., 2016) and Pathview (Luo & Brouwer, 2013) packages based on the relative ion counts and statistical measures given by Metabolon.

### Growth Assays

For assays of growth on alanine as a carbon or nitrogen source, an equal molarity of carbon or nitrogen atoms were added for each indicated molecule to our base minimal media lacking NaNO_3_ and glucose. Agar plates were inoculated with 10^3 conidia and incubated for 72 hours at 37°C 5% CO_2_ in either ambient oxygen or in a chamber that maintained oxygen at a concentration of 0.2% (InvivO_2_ 400 Workstation, Ruskinn Baker). Biofilm biomass cultures were inoculated with 20mL of 10^5 conidia/mL in GMM and grown in petri plates for 24 hours at 37°C 5% CO_2_. Supernatants and air-liquid interface growth were removed, and biofilms were harvested using a cell scraper. Biomass was washed 2X with ddH_2_O with centrifuging at 5000RPM for 10 minutes to spin down biomass, frozen at −80°C, lyophilized, and dry weight was measured. For liquid kinetic growth assays of biofilms, 200μL of 10^5 conidia/mL in GMM was inoculated in six technical replicates per strain in a 96 well plate. Plates were incubated statically in a plate reader at 37°C with Abs_405_ readings every 15 minutes over the first 24 hours of growth.

### Crystal Violet Adherence Assay

U-bottomed 96-well plates were inoculated with 10^5 conidia/mL in GMM and incubated statically for the indicated time at 37°C 5% CO_2_ to allow biofilms to form. To remove non-adherent cells, media was removed, and biofilms were washed twice with water via immersion followed by banging plate onto a stack of paper towels. Adherent biomass was stained with 0.1% (w/v) crystal violet for 10 minutes and biofilms were washed twice with water to remove excess crystal violet. Crystal violet was then dissolved in 100% ethanol, supernatants were transferred to a flat-bottomed plate, and absorbance at 600nm was quantified. Dose-response assays testing the impact of β-chloro-L-alanine on adherence were fit with a non-linear regression based on a dose-response model (GraphPad Prism 9) to calculate EC_50_ values. For assays testing the impact of collagen coating, 50μL of Collagen Coating Solution (Sigma) was applied to half the wells of a 96-well U-bottom plate overnight at room temperature. The solution was removed, and wells were washed one time with PBS prior to inoculation. All crystal violet adherence assays were performed with 3-6 technical replicates and data presented represent at least three biological replicates.

### Oxygen Quantification

Oxygen was quantified as previously described (Kowalski et al., 2020) using a Unixense Oxygen Measuring System 1-CH (Unisense OXY METER) equipped with a micromanipulator (Unisense MM33), motorized micromanipulator stage (Unisense MMS), motor controller (Unisense MC-232), and a 25μm Clark-type/amperometric oxygen sensor (Unisense OX-25). The SensorTrace Suite Software v3.1.151 (Unisense) was utilized to obtain and analyze the data. Falcon 35mm petri dishes (Fisher) were coated with 2mL of 0.6% agar GMM to protect the microelectrode from breaking when performing deep profiling into the biofilms. 3mL of 10^5 conidia/mL in GMM was inoculated into the plates and incubated for 24 hours at 37°C 5% CO_2_. The meniscus of the culture was ∼3mm above the surface of the agar pad, and thus oxygen was measured at the center of each culture in 200μm steps, with technical duplicates at each step, from the air-liquid interface to 2800μm into the culture. Oxygen quantification was performed immediately upon removal of the culture from the incubator. At least seven independent biofilms were measured for each strain across two experiments along with three media only cultures that lacked fungus.

### Fluorescent Microscopy

Fluorescent confocal microscopy was performed on an Andor W1 Spinning Disk Confocal with a Nikon Eclipse Ti inverted microscope stand.

#### AlaA Localization Studies

Af293*alaA-GFP*, Af293*alaA^K322A^-GFP*, and Af293 were cultured in GMM on MatTek^®^ dishes at 37°C in GMM until germlings were visible on an inverted light microscope, ∼9 hours for Af293*alaA-GFP* and Af293, and ∼10 hours for Af293*alaA^K322A^-GFP*. Media was removed and replaced with fresh GMM containing 100nm MitoTracker™ Deep Red FM (ThermoFisher). Cultures were incubated for 30 minutes at 37°C to allow mitochondrial staining. Images were acquired with a 60X oil-immersion objective at 488nm (GFP) and 637nm (MitoTracker) on the Andor W1 Spinning Disk Confocal. Images were deconvolved and max intensity z-projections were generated using the Nikon NIS-Elements AR software. Experiment was also performed with WT Af293 as a negative control for autofluorescence (Figure 2-figure supplement 2). At least ten images for each strain were taken across four replicate cultures.

#### Fungal biofilm imaging and quantification

Biofilms were grown in 2mL of GMM at 10^5 conidia/mL in 35mm glass bottom MatTek^®^ dishes for the indicated duration of time at 37°C 5% CO_2_. At the indicated time, 1mL of media was removed and 500uL of 30μg/mL FITC-SBA (Vector Laboratories) was added to each culture. Cultures were incubated at room temperature for 1 hour to allow staining. Biofilms were then fixed via addition of 500uL 4% paraformaldehyde in PBS and counterstained with 200uL of 275μg/mL calcofluor white (Fluorescent Brightener-28 (Sigma)). Images of the first ∼300μm of the biofilms were acquired on the Andor W1 Spinning Disk Confocal with a 20X multi-immersion objective (Nikon) at 405nm (calcofluor white) and 488nm (FITC-SBA).

For quantification and analysis of biofilms the BiofilmQ framework was used. A detailed explanation of BiofilmQ can be found in previous publications (Hartmann et al., 2021). Briefly, both biomass (calcofluor white) and matrix (FITC-SBA) were thresholded and then segmented into discrete objects with 20 voxel cubes for further analysis. Total biovolume measures were achieved by taking the summed volume of the segmented calcofluor white signal for each biofilm. Total matrix intensity was determined by summing the intensity signal of FITC-SBA in each cube of segmented matrix signal across the entire image. We measured hyphal associated matrix by taking the sum of FITC-SBA intensity that was overlapping in each cube of segmented biomass. Representative images were rendered using the VTK output feature of BiofilmQ. These files could then be rendered in ParaView (Ayachit, 2015) using Ospray ray tracing. Matrix intensity per individual cube as determined in BiofilmQ was then mapped onto the segmented matrix images.

#### Cell wall staining and quantification

Strains were grown in GMM in the center of MatTek^®^ dishes until germlings were visible by an inverted light microscope, ∼9 hours for Af293 and Af293*alaA^rec^* and ∼10 hours for Af293Δ*alaA*. Supernatants were removed and cells were washed with PBS. For calcofluor white staining, germlings were fixed with 4% paraformaldehyde for 15 minutes, washed with PBS and stained with 25μg/mL calcofluor white (Fluorescent Brightener 28, Sigma) in PBS for 15 minutes. Calcofluor white was removed, germlings were washed with PBS, and maintained in 2mL PBS at room-temperature until imaging. For FITC-WGA staining, germlings were stained with 5μg/mL FITC-WGA in GMM for 30 minutes at room temperature. Germlings were washed with PBS and fixed with 4% paraformaldehyde for 15 minutes. WGA stained germlings were then washed with PBS and maintained in 2mL PBS at room temperature until imaging. Finally, for Dectin-1 staining germlings were fixed with 4% paraformaldehyde for 15 minutes and washed with PBS. Blocking solution (RPMI + 10% FCS + 0.025% Tween-20) was applied for 1 hour at room temperature. Blocking solution was removed and 5μg/mL of Dectin-1-Fc in blocking solution was applied for 1 hour at room temperature. Germlings were washed with PBS and the secondary antibody AlexaFluor 488 anti-human IgG (ThermoFisher) was added at a 1/300 dilution in PBS for 1 hour at room temperature. Germlings were washed one final time with PBS and cells were maintained in 2mL PBS until imaging. All staining took place in the dark and extreme care was taken to not disrupt the Af293Δ*alaA* germlings.

All germlings were imaged on the Andor W1 Spinning Disk Confocal with a 60X oil-immersion objective using 405nm for calcofluor white and 488nm for FITC-WGA and Dectin-1-Fc. Cell wall staining was quantified using Fiji (ImageJ). Z-stacks were assembled using a sum-intensity Z-projection. Regions of interest (ROI’s) were drawn around each individual germling within a given image, along with a region lacking any germlings to account for background fluorescence. Within each ROI the area, sum intensity, and mean intensity were quantified. To obtain corrected mean intensity measurements, the mean background intensity was multiplied by the area of the ROI to calculate total background contribution. The total background contribution was subtracted from the ROI’s sum intensity and this value was divided by the area of the ROI yielding the final corrected mean fluorescence intensity. Each cell wall stain was performed in triplicate cultures and at least three fields of view were obtained for each culture. For FITC-WGA the staining pattern was almost entirely absent from the germ-tube and enough natural size heterogeneity was found both within and between cultures to act as a confounding variable. Thus, for FITC-WGA ROIs were drawn around each conidial body, where the staining was present, rather than the entire germling.

### Extracellular matrix monosaccharide analysis and ELLA

#### Enzyme Linked Lectin Assay (ELLA)

100µL of 10^5 conidia per mL in GMM were inoculated into wells of 96 well plate and incubated for 24h. Culture supernatants were then transferred to a 384 well plate Immulon 4HBX with or without 500pM of recombinant Agd3. After a 1-hour incubation period, wells were washed three times with 1X TBS – 0.05% Tween20. A preincubated solution of 30nM soybean agglutinin lectin coupled to biotin and 1/700 avidin-HRP in TBS-T was added to the wells and incubated for 1 hour. After 3 TBS-T washes, detection was performed using Ultrasensitive TMB read at 450nm. Normalization of the values were performed reporting the absorbance reads to the absorbance of Af293.

#### Extracellular Matrix Monosaccharide Composition by Gas Chromatography Coupled to Mass Spectrometry

100ml of GMM was inoculated with 10^4 conidia per mL and incubated for 3 days at 37°C at 200rpm. Culture supernatants were filtered by Miracloth prior to being dialyzed for 3 days against Milli-Q^®^ water and lyophilized. About 0.5mg of dried material was then derivatized into TriMethylSilyl derivatives. Samples were hydrolyzed with either 2 M trifluoroacetic acid for 2 hours at 110°C or 6 M hydrochloric acid (HCl) for 4 hours at 100°C. Monosaccharides were then converted in methyl glycosides by heating in 1 M methanol-HCl (Sigma-Aldrich) for 16 hours at 80°C. Samples were dried and washed twice with methanol prior to re-N-acetylating hexosamine residues. Re-N-acetylation was performed by incubation with a mix of methanol, pyridine, acetic anhydride (10:2:3) for 1 hour at room temperature. Samples were then treated with hexamethyldisilazane-trimethylchlorosilane-pyridine solution (3:1:9; ThermoFisher) for 20 min at 110°C. The resulting TMS methyl glycosides were dried, resuspended in 1 ml of cyclohexane, and injected in the Agilent 7890B GC-5977A MSD. Identification and quantification of the monosaccharides was performed using a mix of monosaccharide calibrants injected at different concentrations as a reference. Quantification was finally normalized to an equivalent of 1mg of material before comparison between groups.

### RNA Extraction and RTqPCR

RNA was extracted from 24-hour biofilm cultures in a 6-well plate. Supernatant was removed and 500μL of TRIsure^TM^ (Bioline Reagents) was immediately applied to the biofilms. Biofilm suspensions were centrifuged, and supernatant was removed. Biomass was resuspended in 200μL TRIsure^TM^ flash frozen in liquid nitrogen, and bead beat with 2.3mm beads. Homogenate was brought to a final volume of 1mL with TRIsure^TM^, bead beaten a second time, and RNA was extracted following the manufacturer’s protocol. 5μg of RNA was DNase treated with TURBO DNA-*free*^TM^ kit (Invitrogen) according to manufacturer’s protocol. 500ng of DNase-treated RNA was run on an agarose gel to ensure RNA integrity. 500ng of DNase-treated RNA was used for cDNA synthesis as previously described (Beattie et al., 2017). The RTqPCR data were collected on a CFX Connect Real-Time PCR Detection System (Bio-Rad) with CFX Maestro Software (Bio-Rad). Gene expression was normalized to *tefA* expression for all experiments. Primers used for RTqPCR are listed in Table S2.

### Antifungal drug susceptibility

#### Biofilm Assays

To test susceptibility of biofilms to inhibition by calcofluor white and caspofungin, 500μL of 10^5 conidia per mL in GMM were inoculated into wells of 24 well plates and biofilms were grown statically for 24 hours at 37°C 5% CO_2_. Any air-liquid interface growth was removed using a sterile pipette tip, the supernatant was removed from each well, and fresh media containing the indicated concentration of calcofluor white or caspofungin was added to the biofilm. Biofilms were incubated for a further 3 hours at 37°C 5% CO_2_, washed with PBS, and 300μL of XTT solution was added to each well (0.5mg/mL XTT [2,3-bis-(2-methoxy-4-nitro-5-sulfophenyl)-2H-tetrazolium-5-carboxanilide] (VWR) with 25μM menadione in PBS). XTT solution was incubated for 1 hour at 37°C to allow reduction of the dye. 150μL of supernatants were transferred to a flat-bottomed 96 well plate and the absorbance at 450nm was read on a plate reader. Abs_450nm_ values of the treated samples were compared to untreated biofilms to calculate the relative metabolic activity as the percent of untreated control for each strain. All XTT assays were performed on at least three biological replicates.

For the adenylate kinase release assay biofilms were grown and treated in the same manner as above. After three hours of echinocandin treatment supernatants were collected and allowed to cool to room temperature. XTT assay was performed on the biofilms according to the protocol above for matched XTT data. Relative adenylate kinase levels were measured on 40μL of supernatants via the ToxiLight^TM^ Non-Destructive Cytotoxicity BioAssay Kit (Lonza) according to the manufacturer’s instructions. Chemiluminescence was measured on a Synergy Neo2 multi-mode plate reader (BioTek). Experiment was performed on four independent biological replicates.

#### Conidia Assays

Minimal effective concentration (MEC) assays were performed by inoculating 100μL of 2*10^5 conidia/mL in GMM into 96 well flat-bottomed plates containing 100μL serial 2-fold dilutions of caspofungin in GMM from 8μg/mL to 0.015625μg/mL along with a no drug control. Cultures were grown statically for 24 hours and viewed under an inverted light microscope for the concentration at which gross morphological changes characteristic of caspofungin treatment became visible. This concentration was deemed the MEC. Radial growth assays were performed by inoculating GMM agar plates containing the indicated quantity of caspofungin with 10^3 conidia in 2μL 0.01% Tween-80 and incubating at 37°C 5% CO_2_ for 72 hours. Images are representative of four biological replicates.

### Murine Fungal Burden Assay

All mice were housed in autoclaved cages at 3-4 mice per cage and provided food and autoclaved water *ad libitum*. Female outbred CD-1 (Charles River Laboratory), 20-24 grams, were immune-suppressed with 150mg/kg cyclophosphamide (Ingenus Pharmaceuticals, LLC) interperitoneally 48 hours prior to inoculation and 40mg/kg triamcinolone acetonide (Kenalog-10, Bristol-Myers Squibb) subcutaneously 24 hours prior to fungal challenge. Mice were administered 10^6 conidia in 30μL PBS intranasally under isoflurane anesthesia. Mock mice were administered 30μL sterile PBS. Micafungin treated mice were administered 1mg/kg micafungin (Mycamine^®^, Astellas Pharma) interperitoneally at 24, 48, and 72 hours post-fungal inoculation. Untreated mice were administered 100μL 0.9% saline (vehicle control) interperitoneally, at 24, 48, and 72 hours post-fungal inoculation. Mice were sacrificed at 84 hours post-fungal inoculation and lungs were harvested for fungal burden.

Lungs were divided between two 2mL screw cap tubes and physically chopped using a dissecting scissors, flash frozen in liquid nitrogen, and lyophilized for 48 hours. The freeze-dried lungs were then bead beaten with 2.3mm Zirconia beads and DNA was extracted using the E.Z.N.A.^®^ Fungal DNA Mini Kit (Omega Bio-tek) with the following modifications. Bead beaten lungs were resuspended in 600μL FG1 buffer, bead beaten a second time and incubated at 65°C for 1 hour. Samples were centrifuged and supernatants from the split lung samples were combined in a new tube. The protocol was continued with 200μL of the combined supernatant according to the manufacturer’s instructions with two elution steps using 100μL molecular grade water heated to 65°C. qPCR quantification of fungal DNA was performed as previously described (Li et al., 2011). The fungal burden experiment was performed two times with n ≥ 6 in each experimental group per experiment and n = 5 mock across the two experiments. Four mice across the two experiments, including one in the Af293Δ*alaA* treated group, were censored for either unsuccessful infection or fungal DNA extraction based on the criteria of having less fungal DNA than the highest mock control value.

## Statistics and Reproducibility

All statistical analyses were performed in GraphPad Prism 9 with the exception of metabolomics statistics which were performed by Metabolon. Unless otherwise noted all experiments were performed with a minimum of three biologically independent samples.

## Acknowledgements

We thank A. Lavanway (Dartmouth) for their microscopy expertise. We also thank the Imaging Facility at Dartmouth and the Biomolecular Targeting Core (P20-GM113132) for use of equipment. This work was supported by funding from the NIH National Institute of Allergy and Infectious Diseases (NIAID) (grant no. R01AI130128 and R01AI146121), a pilot award from the Cystic Fibrosis Foundation (CFF) Research Development Award (STANTO15RO), and a CFF research award (CRAMERGO19). J.K. was supported by the Molecular Pathogenesis Training Grant (T32AI007519).

## Supplemental Figures and Tables

**Figure 1-figure supplement 1:**
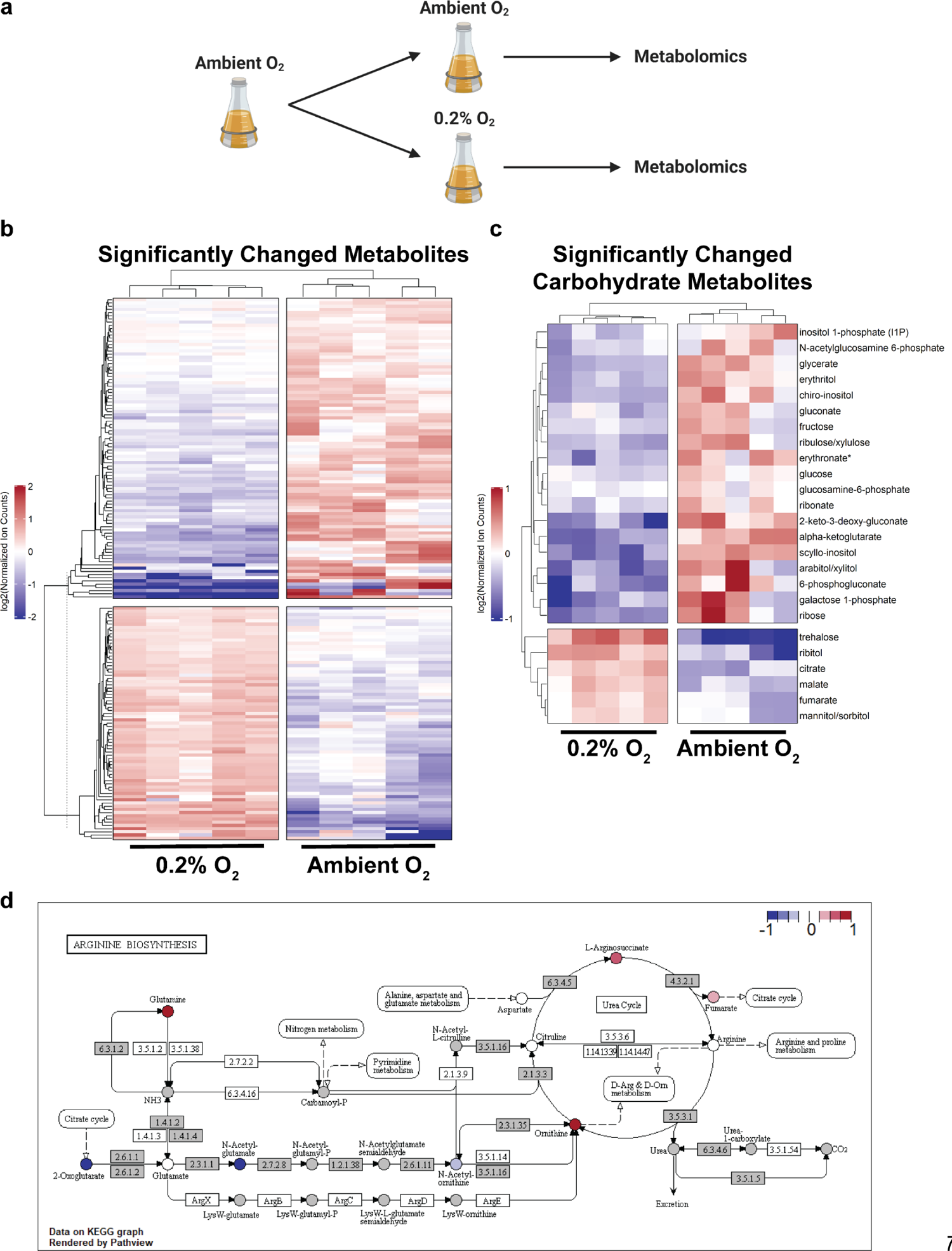
Acute exposure to a low oxygen environment significantly changes *A. fumigatus* metabolism. A) Outline of the metabolomics experiment. Shaking culture were grown for 24 hours in liquid GMM followed by either a shift to a 0.2% oxygen environment or continued incubation in ambient oxygen for two hours. Biomass was then harvested, flash frozen to quench metabolic reactions, lyophilized, and submitted for metabolomics. B) Overview of all significantly altered metabolites detected in experiment. C) Significantly changed carbohydrate related metabolites in the two conditions. For C-D ion counts were normalized to the average ion count for the respective metabolite across all samples and log_2_ transformed. Each column corresponds to an individual sample (n = 5 per condition). D) Average relative abundance of metabolites in 0.2% vs ambient oxygen (log_2_ transformed) mapped onto the Arginine Biosynthesis KEGG map using the Pathview R package. Red indicates greater abundance in 0.2% oxygen, blue indicates greater abundance in ambient oxygen, white indicates no difference between conditions, and metabolites not detected are in grey.

**Figure 1-figure supplement 2:**
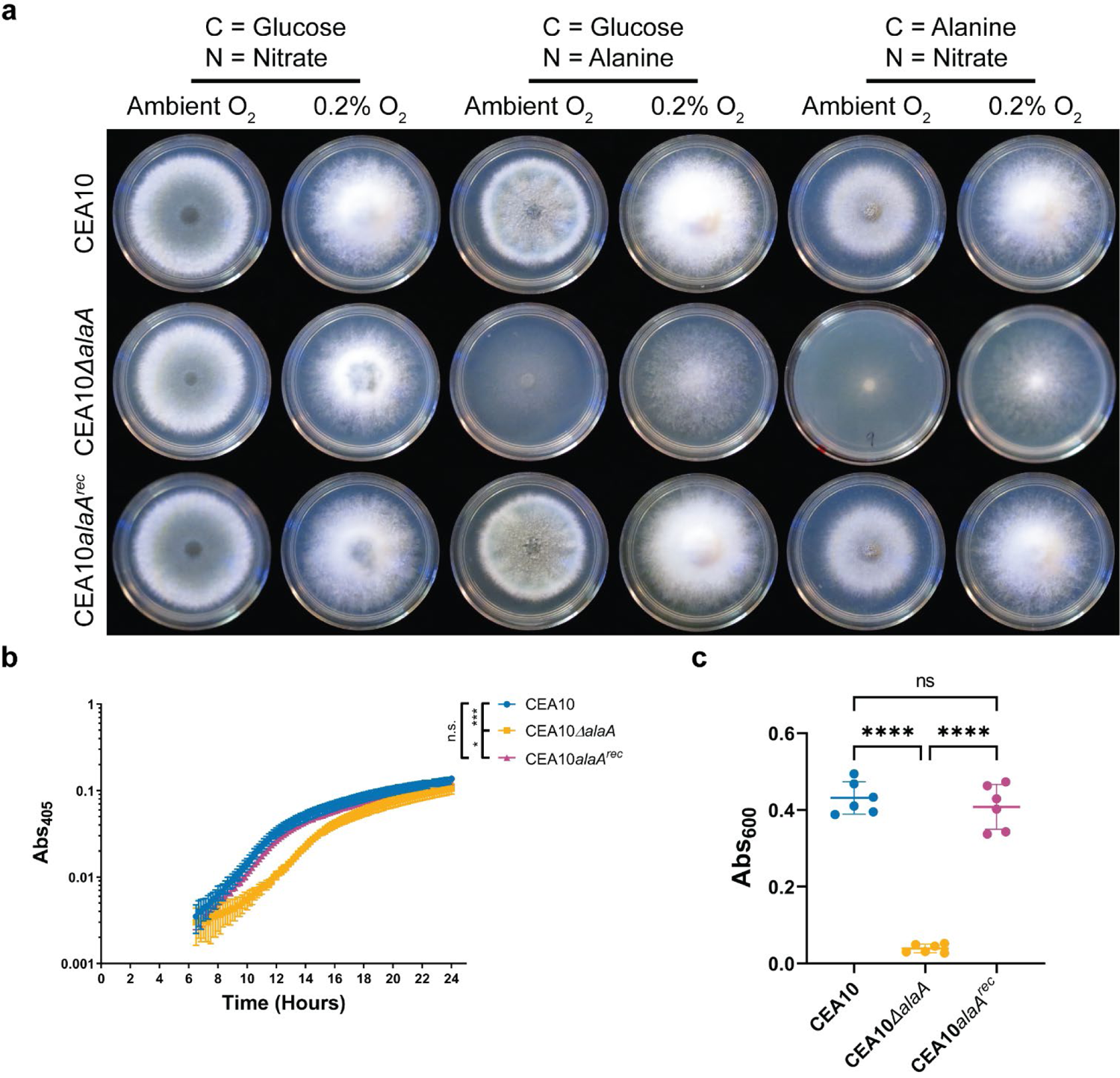
*alaA* is required for alanine catabolism and biofilm physiology in the CEA10 strain background. A) Growth of CEA10Δ*alaA* on minimal media containing the indicated sole carbon and nitrogen sources in ambient oxygen and 0.2% oxygen environments. B) Static growth assay of CEA10Δ*alaA* over the first 24 hours of biofilm growth. Mean +/- SD of 6 technical replicates is shown. Experiment was repeated a minimum of 3 times with similar results. * p < 0.05, *** p < 0.001 by One-Way ANOVA with a Tukey’s multiple comparisons test C) Crystal violet adherence assay of 24-hour biofilms (n = 6). **** p < 0.0001, n.s. = not significant as determined by One-Way ANOVA with a Tukey’s multiple comparisons test.

**Figure 1-figure supplement 3:**
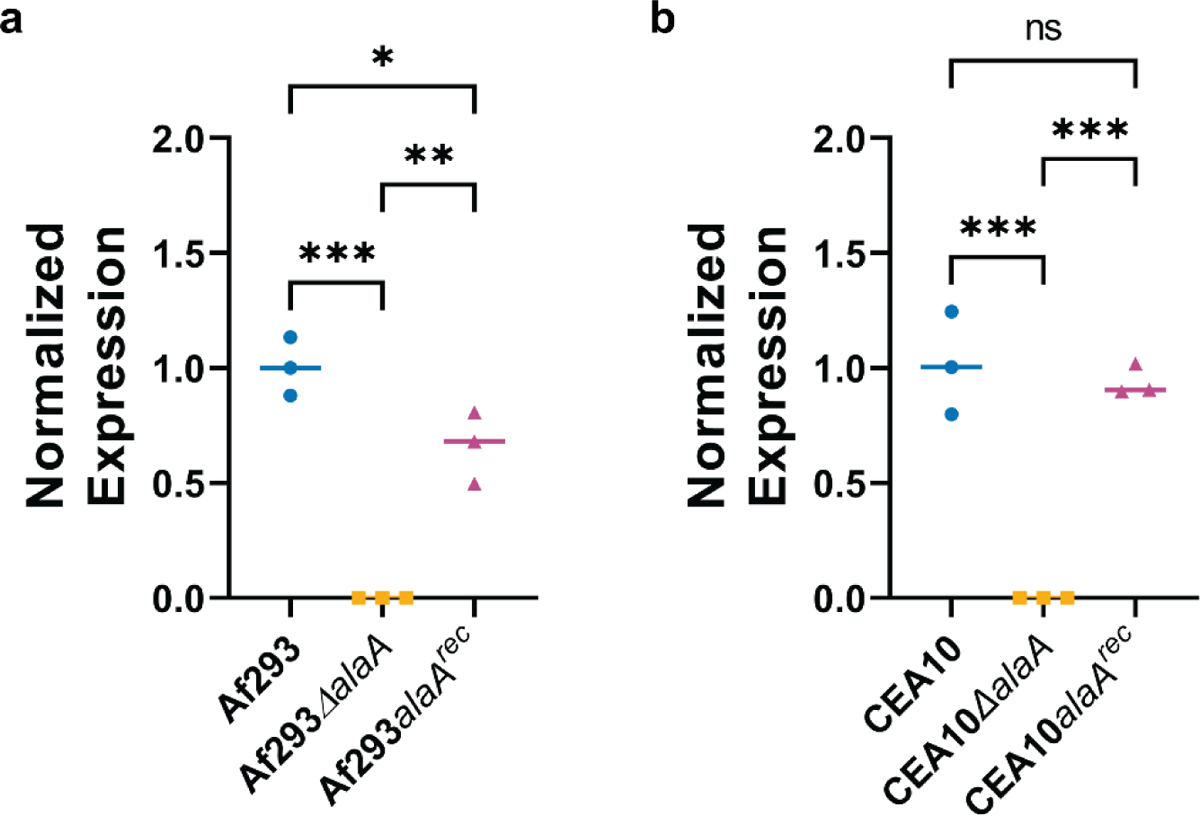
RTqPCR of *alaA* expression in *alaA* deletion and reconstituted strains. A-B) RNA was harvested from 24-hour biofilms and *alaA* expression was determined by RTqPCR (n = 3). Each replicate along with the median are shown. * p < 0.05, ** p < 0.01, *** p < 0.001, n.s. = not significant as determined by One-Way ANOVA with a Tukey’s multiple comparisons test.

**Figure 2-figure supplement 1:**
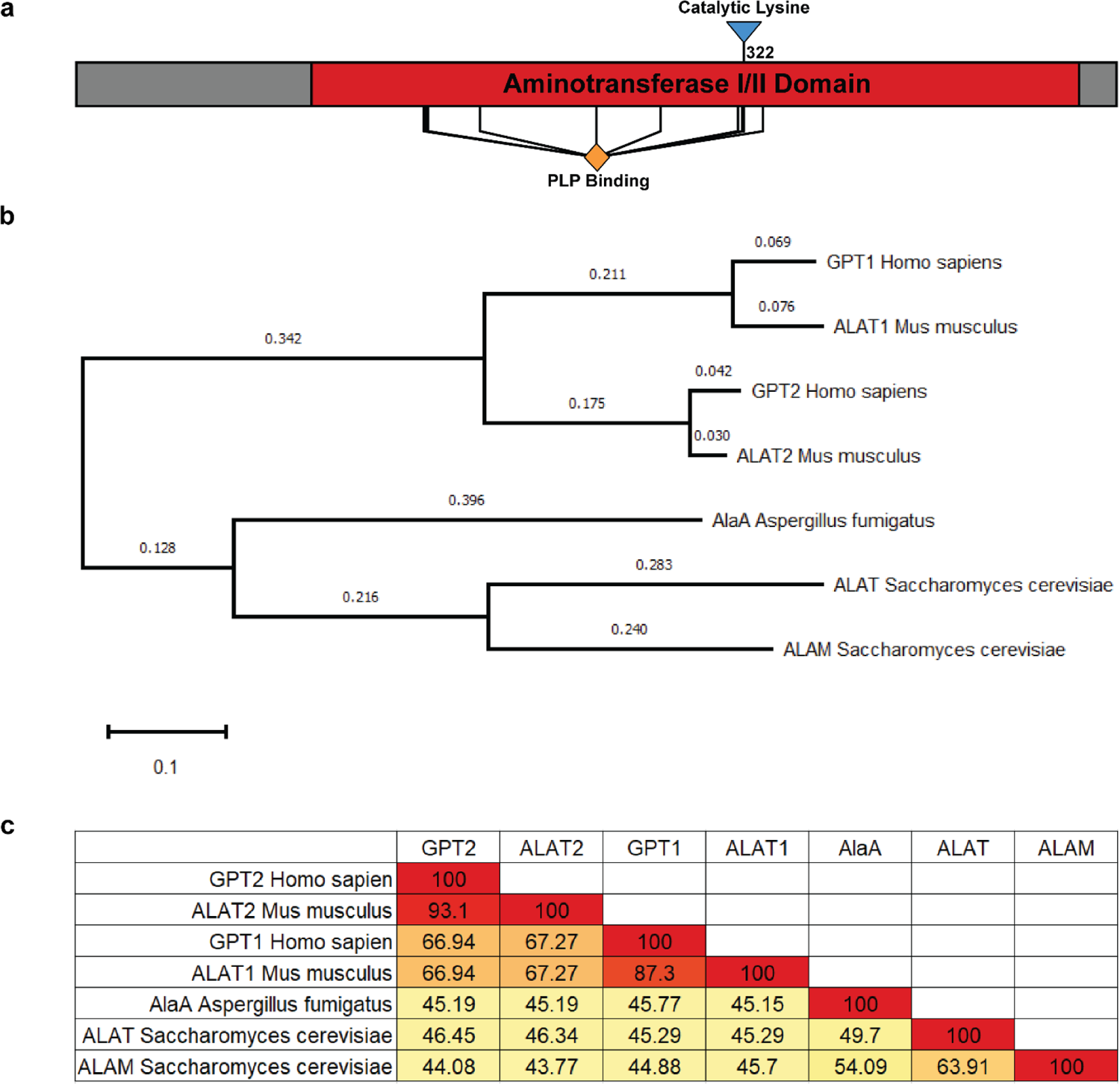
AlaA protein domain architecture and degree of similarity to alanine aminotransferases of several model systems. A) AlaA linear protein structure highlighting the aminotransferase class I/II domain, predicted PLP-binding residues, and the catalytic lysine residue mutated in the catalytic null strain (Af293*alaA^K322A^-GFP*). B) Phylogeny of AlaA relative to human, murine, and *Saccharomyces cerevisiae* alanine aminotransferases. All species except *A. fumigatus* encode two alanine aminotransferases in their genomes. Proteins were aligned in MEGA X using MUSCLE and a maximum-likelihood tree was generated. Scale bar and branch lengths refer to substitutions per site. C) Percent identity matrix of the alanine aminotransferases shown in (B).

**Figure 2-figure supplement 2:**
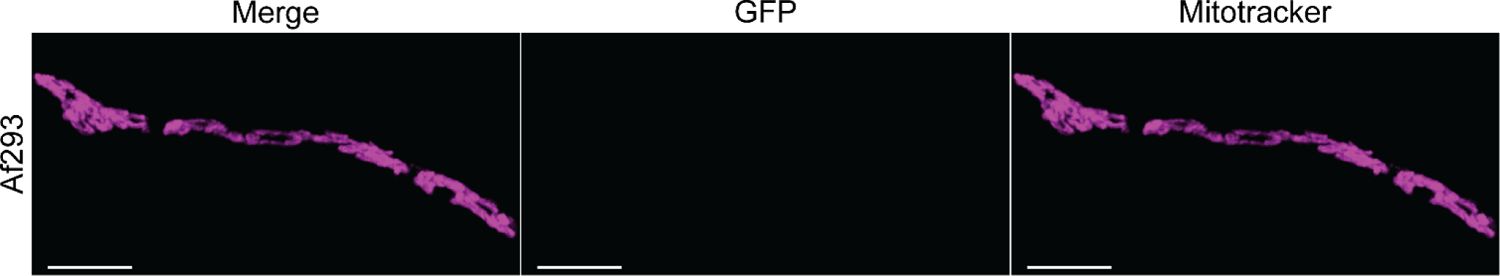
Af293 wildtype control for AlaA localization experiments. Representative micrographs of wildtype Af293 germlings stained with Mitotracker^TM^ Deep Red FM (magenta). No fluorescence was observed in the GFP channel.

**Figure 5-figure supplement 1:**
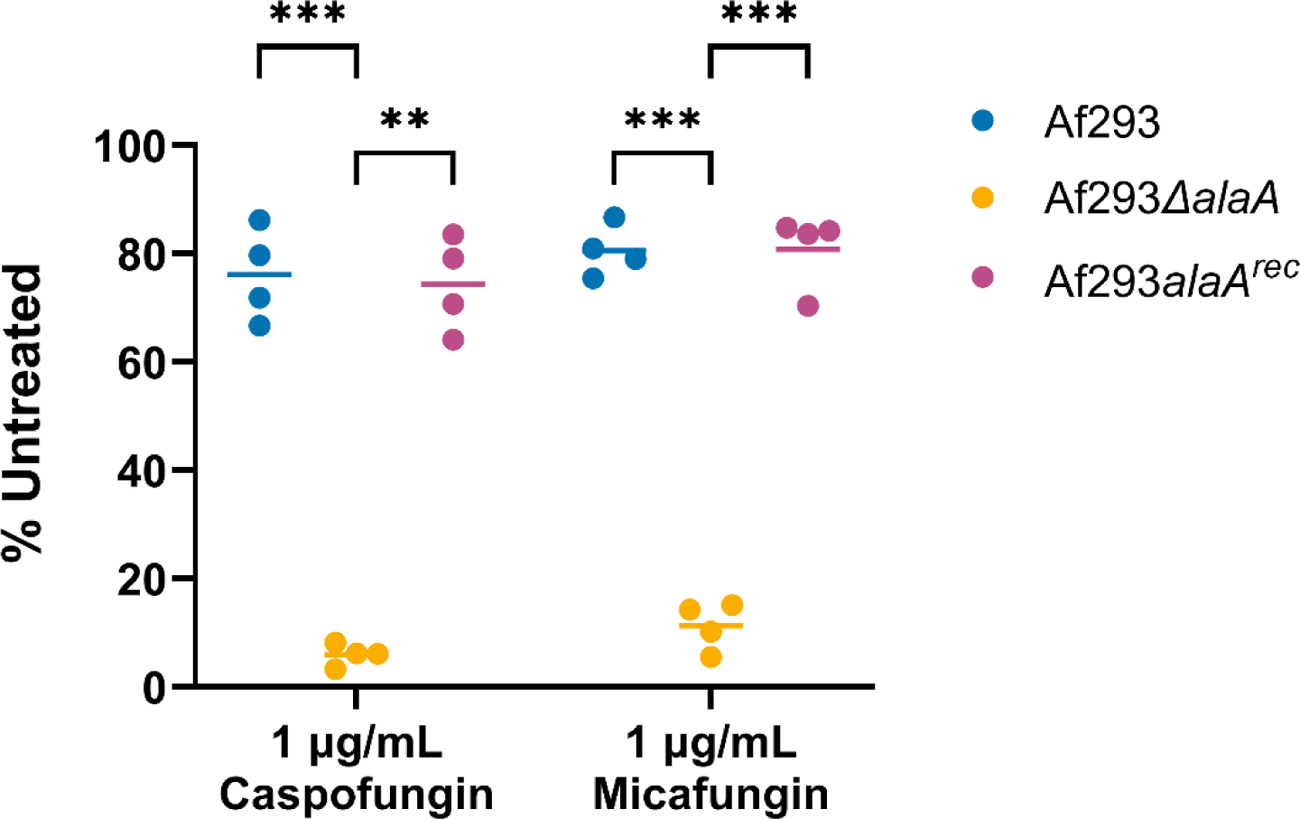
XTT assay corresponding to the cultures used in the adenylate kinase release assay. Biofilms were grown for 24 hours and treated with 1µg/mL caspofungin (left) or micafungin (right) for 3 hours. Supernatants were used to quantify adenylate kinase activity (Figure 5I) and an XTT assay was performed to measure viability of biofilm biomass. Each replicate and mean are shown (n = 4). ** p < 0.01, *** p < 0.001 as determined by Two-Way ANOVA with a Tukey’s multiple comparisons test.

**Figure 5-figure supplement 2:**
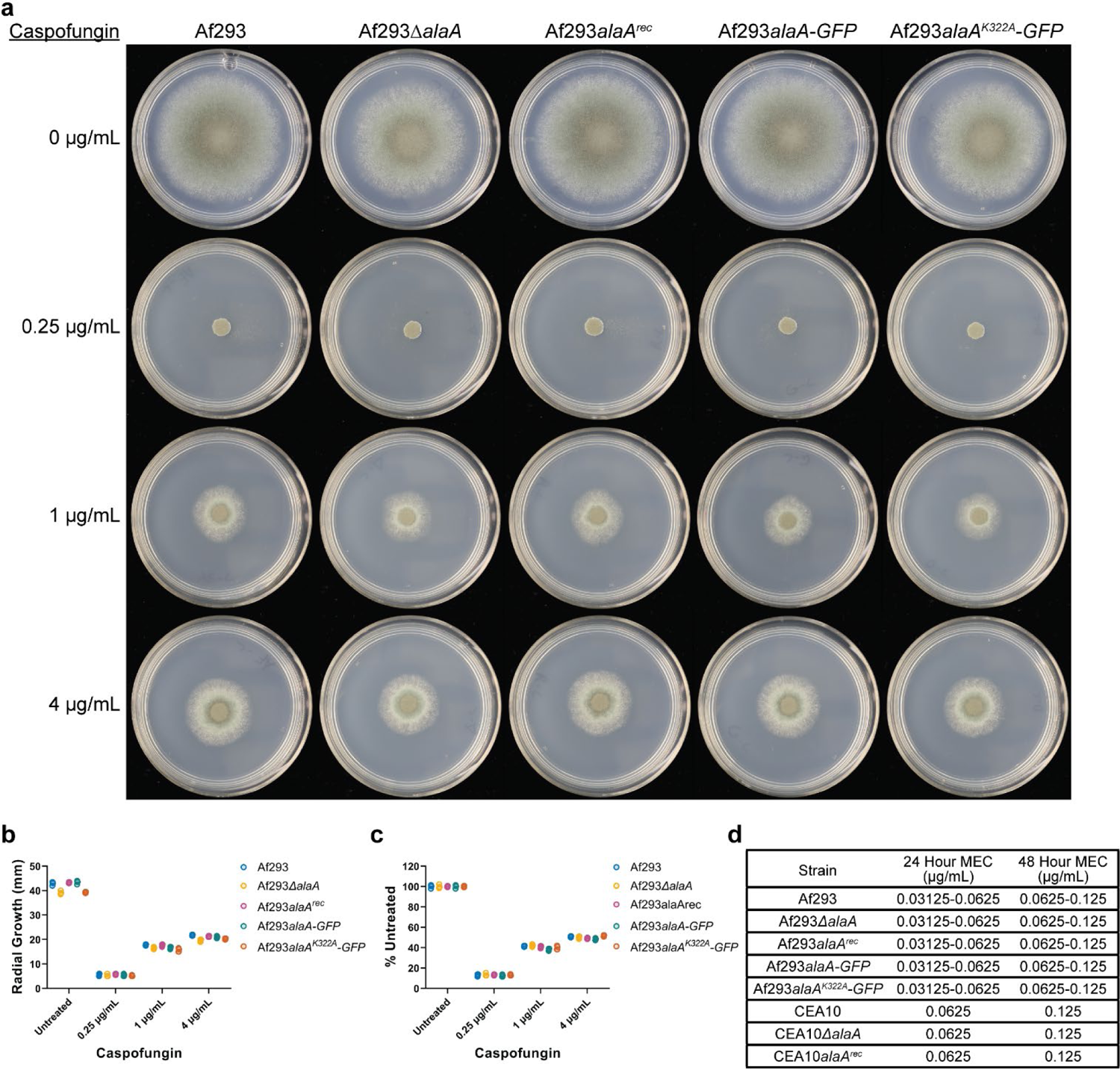
Increased susceptibility of *alaA* null strains to caspofungin is biofilm specific. A) Conidial radial growth assays of the indicated strains grown on the indicated concentrations of caspofungin in GMM for 72 hours. Images are representative of four replicate cultures. B) Quantification of radial growth as the mm diameter for each colony (n = 4). C) Radial growth normalized to the untreated control for each strain. Individual replicates and mean are shown (n = 4) for (B-C). D) Minimum effective concentration (MEC) of caspofungin for the indicated strains at 24 and 48 hours of incubation in GMM containing increasing concentrations of caspofungin (n = 3).

**Figure 6-figure supplement 1:**
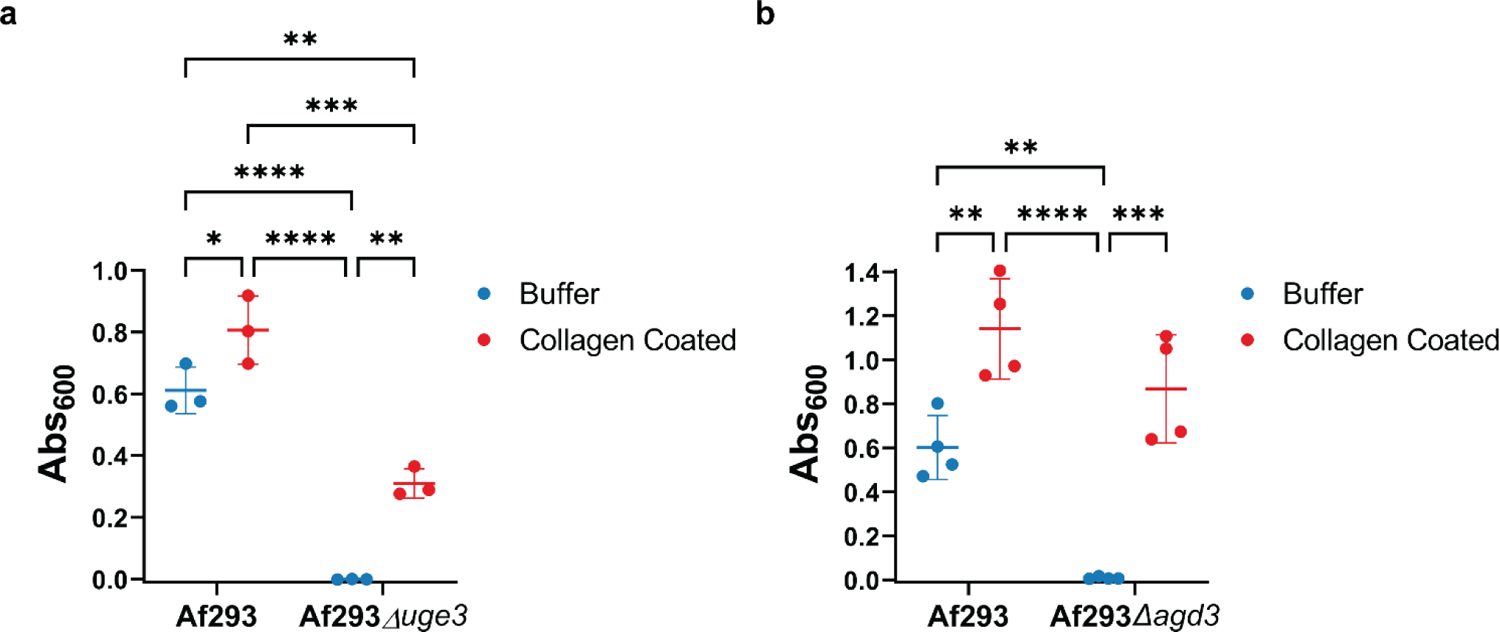
Collagen coating partially rescues adherence of Af293Δ*uge3* and Af293Δ*agd3*. A-B) Crystal violet adherence assay of Af293Δ*uge3* (A) and Af293Δ*agd3* (B) in wells of a 96-well plate coated with collagen or PBS (buffer). Each replicate along with the mean +/- SD are shown (n ≥ 3). * p < 0.05, ** p < 0.01, *** p < 0.001, **** p < 0.0001 as determined by Two-Way ANOVA with a Tukey’s multiple comparisons test.

**Table S1:**
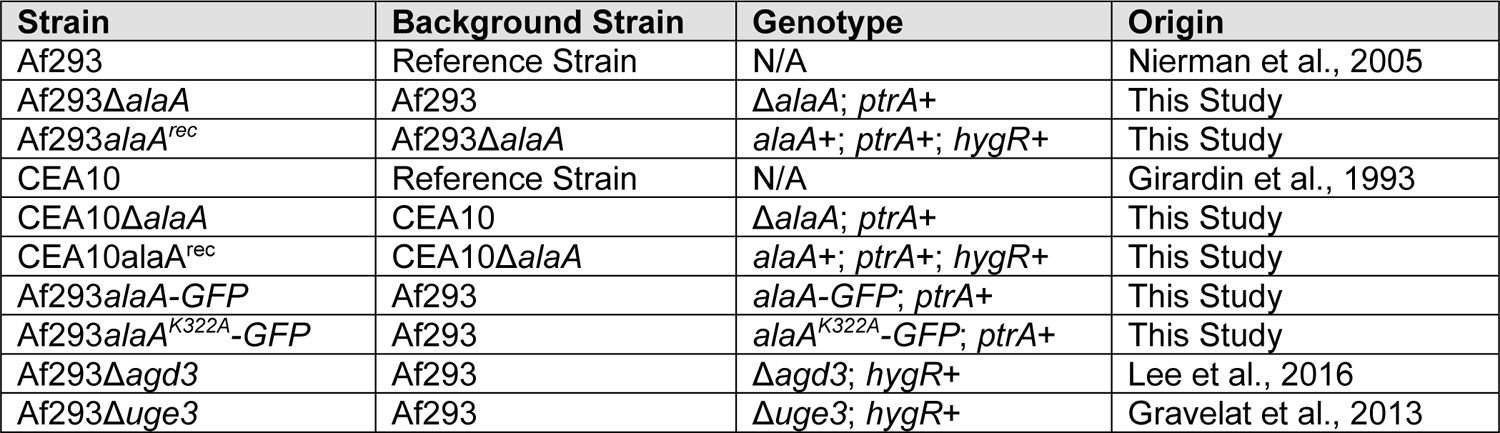
Fungal strains used in this study.

**Table S2:**
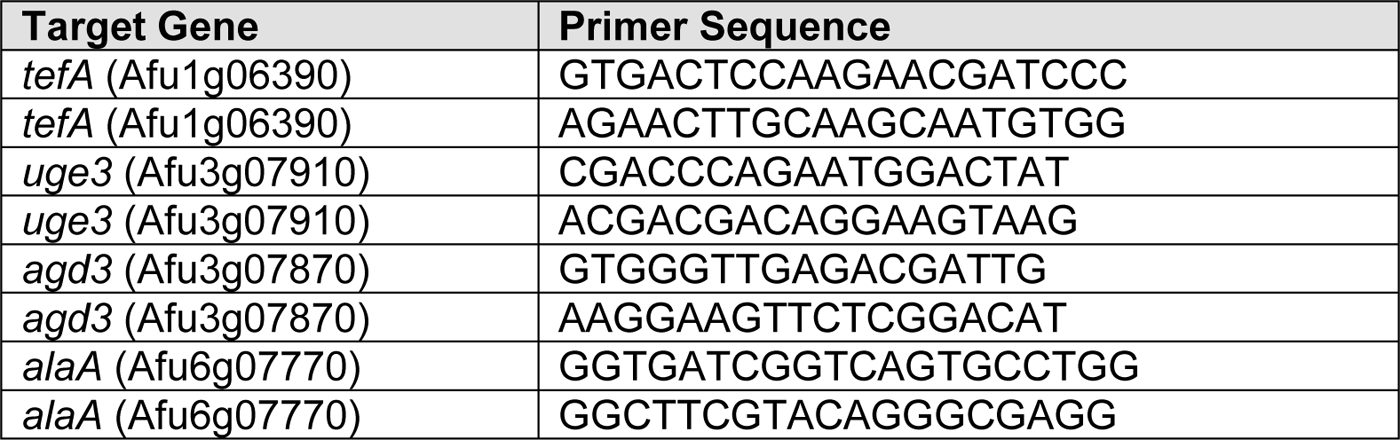
**Primers used for RTqPCR.**

## References

1. Aruanno, M., Glampedakis, E., & Lamoth, F. (2019). Echinocandins for the treatment of invasive aspergillosis: From laboratory to bedside. In Antimicrobial Agents and Chemotherapy (Vol. 63, Issue 8). American Society for Microbiology. https://doi.org/10.1128/AAC.00399-19

2. Ayachit, U. (2015). The paraview guide (full color version): A parallel visualization application. Kitware.

3. Bamford, N. C., Le Mauff, F., Van Loon, J. C., Ostapska, H., Snarr, B. D., Zhang, Y., Kitova, E. N., Klassen, J. S., Codée, J. D. C., Sheppard, D. C., & Howell, P. L. (2020). Structural and biochemical characterization of the exopolysaccharide deacetylase Agd3 required for Aspergillus fumigatus biofilm formation. Nature Communications, 11(1), 1–13. https://doi.org/10.1038/s41467-020-16144-5

4. Barker, B. M., Kroll, K., Vödisch, M., Mazurie, A., Kniemeyer, O., & Cramer, R. A. (2012). Transcriptomic and proteomic analyses of the *Aspergillus fumigatus* hypoxia response using an oxygen-controlled fermenter. BMC Genomics, 13(1), 62. https://doi.org/10.1186/1471-2164-13-62

5. Beattie, S. R., Mark, K. M. K. K., Thammahong, A., Nicolas, L., Ries, A., Dhingra, S., Caffrey-Carr, A. K., Cheng, C., Black, C. C., Bowyer, P., Bromley, M. J., Obar, J. J., Goldman, G. H., Cramer, R. A., Ries, L. N. A., Dhingra, S., Caffrey-Carr, A. K., Cheng, C., Black, C. C., … Cramer, R. A. (2017). Filamentous fungal carbon catabolite repression supports metabolic plasticity and stress responses essential for disease progression. PLoS Pathogens, 13(4), 1–29. https://doi.org/10.1371/journal.ppat.1006340

6. Beuster, G., Zarse, K., Kaleta, C., Thierbach, R., Kiehntopf, M., Steinberg, P., Schuster, S., & Ristow, M. (2011). Inhibition of alanine aminotransferase in Silico and in vivo promotes mitochondrial metabolism to impair malignant growth. Journal of Biological Chemistry, 286(25), 22323–22330. https://doi.org/10.1074/jbc.M110.205229

7. Bhabhra, R., & Askew, D. S. (2005). Thermotolerance and virulence of Aspergillus fumigatus: Role of the fungal nucleolus. In Medical Mycology (Vol. 43, Issue SUPPL.1, pp. S87–S93). Oxford Academic. https://doi.org/10.1080/13693780400029486

8. Casadevall, A. (2017). The Pathogenic Potential of a Microbe. MSphere, 2(1). https://doi.org/10.1128/msphere.00015-17

9. Chen, Y., Mauff F. Le, Wang, Y., Lu, R., Sheppard, D. C., Lu, L., & Zhang, S. (2020). The transcription factor soma synchronously regulates biofilm formation and cell wall homeostasis in aspergillus fumigatus. MBio, 11(6), 1–16. https://doi.org/10.1128/mBio.02329-20

10. Chung, D., Barker, B. M., Carey, C. C., Merriman, B., Werner, E. R., Lechner, B. E., Dhingra, S., Cheng, C., Xu, W., Blosser, S. J., Morohashi, K., Mazurie, A., Mitchell, T. K., Haas, H., Mitchell, A. P., & Cramer, R. A. (2014). ChIP-seq and In Vivo Transcriptome Analyses of the Aspergillus fumigatus SREBP SrbA Reveals a New Regulator of the Fungal Hypoxia Response and Virulence. PLoS Pathogens, 10(11), e1004487. https://doi.org/10.1371/journal.ppat.1004487

11. Di Piazza, S., Houbraken, J., Meijer, M., Cecchi, G., Kraak, B., Rosa, E., & Zotti, M. (2020). Thermotolerant and thermophilic mycobiota in different steps of compost maturation. Microorganisms, 8(6), 1–9. https://doi.org/10.3390/microorganisms8060880

12. Didone, L., Oga, D., & Krysan, D. J. (2011). A novel assay of biofilm antifungal activity reveals that amphotericin B and caspofungin lyse Candida albicans cells in biofilms. Yeast, 28(8), 561–568. https://doi.org/10.1002/yea.1860

13. Feala, J. D., Coquin, L., McCulloch, A. D., & Paternostro, G. (2007). Flexibility in energy metabolism supports hypoxia tolerance in Drosophila flight muscle: Metabolomic and computational systems analysis. Molecular Systems Biology, 3(1), 99. https://doi.org/10.1038/msb4100139

14. Felig, P. (1973). The glucose-alanine cycle. Metabolism, 22(2), 179–207. https://doi.org/10.1016/0026-0495(73)90269-2

15. Felig, P., Pozefsk, T., Marlis, E., & Cahill, G. F. (1970). Alanine: Key role in gluconeogenesis. Science, 167(3920), 1003–1004. https://doi.org/10.1126/science.167.3920.1003

16. Fontaine, T., Delangle, A., Simenel, C., Coddeville, B., van Vliet, S. J., van Kooyk, Y., Bozza, S., Moretti, S., Schwarz, F., Trichot, C., Aebi, M., Delepierre, M., Elbim, C., Romani, L., & Latgé, J. P. (2011). Galactosaminogalactan, a new immunosuppressive polysaccharide of Aspergillus fumigatus. PLoS Pathogens, 7(11), 1002372. https://doi.org/10.1371/journal.ppat.1002372

17. Girardin, H., Latge, J. P., Srikantha, T., Morrow, B., & Soll, D. R. (1993). Development of DNA probes for fingerprinting Aspergillus fumigatus. Journal of Clinical Microbiology, 31(6), 1547–1554. https://doi.org/10.1128/jcm.31.6.1547-1554.1993

18. Golichowski, A., & Jenkins, W. T. (1978). Inactivation of Pig Heart Alanine Aminotransferase by P-Chloroalanine’. In AIKHIVES OF BIOCHEMISTI∼Y ANI) BI∼IWYSI∼∼S (Vol. 189, Issue 1).

19. Grahl, N., Puttikamonkul, S., Macdonald, J. M., Gamcsik, M. P., Ngo, L. Y., Hohl, T. M., & Cramer, R. A. (2011). In vivo hypoxia and a fungal alcohol dehydrogenase influence the pathogenesis of invasive pulmonary aspergillosis. PLoS Pathogens, 7(7). https://doi.org/10.1371/journal.ppat.1002145

20. Gravelat, F. N., Beauvais, A., Liu, H., Lee, M. J., Snarr, B. D., Chen, D., Xu, W., Kravtsov, I., Hoareau, C. M. Q. Q., Vanier, G., Urb, M., Campoli, P., Al Abdallah, Q., Lehoux, M., Chabot, J. C., Ouimet, M. C., Baptista, S. D., Fritz, J. H., Nierman, W. C., … Sheppard, D. C. (2013). Aspergillus Galactosaminogalactan Mediates Adherence to Host Constituents and Conceals Hyphal β-Glucan from the Immune System. PLoS Pathogens, 9(8), e1003575. https://doi.org/10.1371/journal.ppat.1003575

21. Gressler, M., Heddergott, C., N’Go, I. C., Renga, G., Oikonomou, V., Moretti, S., Coddeville, B., Gaifem, J., Silvestre, R., Romani, L., Latgé, J. P., & Fontaine, T. (2019). Definition of the Anti-inflammatory Oligosaccharides Derived From the Galactosaminogalactan (GAG) From Aspergillus fumigatus. Frontiers in Cellular and Infection Microbiology, 9, 365. https://doi.org/10.3389/fcimb.2019.00365

22. Gu, Z., Eils, R., & Schlesner, M. (2016). Complex heatmaps reveal patterns and correlations in multidimensional genomic data. Bioinformatics, 32(18), 2847–2849. https://doi.org/10.1093/bioinformatics/btw313

23. Gugnani, H. C. (2003). Ecology and taxonomy of pathogenic aspergilli. In Frontiers in Bioscience (Vol. 8, Issue SUPPL., pp. 346–357). Frontiers in Bioscience. https://doi.org/10.2741/1002

24. Hagiwara, D., Miura, D., Shimizu, K., Paul, S., Ohba, A., Gonoi, T., Watanabe, A., Kamei, K., Shintani, T., Moye-Rowley, W. S., Kawamoto, S., & Gomi, K. (2017). A Novel Zn2-Cys6Transcription Factor AtrR Plays a Key Role in an Azole Resistance Mechanism of Aspergillus fumigatus by Co-regulating cyp51A and cdr1B Expressions. PLoS Pathogens, 13(1), e1006096. https://doi.org/10.1371/journal.ppat.1006096

25. Harrison, J. F. (2015). Handling and use of oxygen by pancrustaceans: Conserved patterns and the evolution of respiratory structures. Integrative and Comparative Biology, 55(5), 802–815. https://doi.org/10.1093/icb/icv055

26. Hartmann, R., Jeckel, H., Jelli, E., Singh, P. K., Vaidya, S., Bayer, M., Rode, D. K. H., Vidakovic, L., Díaz-Pascual, F., Fong, J. C. N., Dragoš, A., Lamprecht, O., Thöming, J. G., Netter, N., Häussler, S., Nadell, C. D., Sourjik, V., Kovács, Á. T., Yildiz, F. H., & Drescher, K. (2021). Quantitative image analysis of microbial communities with BiofilmQ. Nature Microbiology, 6(2), 151–156. https://doi.org/10.1038/s41564-020-00817-4

27. Herbrecht, R., Patterson, T. F., Slavin, M. A., Marchetti, O., Maertens, J., Johnson, E. M., Schlamm, H. T., Donnelly, J. P., & Pappas, P. G. (2015). Application of the 2008 definitions for invasive fungal diseases to the trial comparing voriconazole versus amphotericin B for therapy of invasive aspergillosis: A Collaborative Study of the Mycoses Study Group (MSG 05) and the European Organization for Research and Treatment of Cancer Infectious Diseases Group. Clinical Infectious Diseases, 60(5), 713–720. https://doi.org/10.1093/cid/ciu911

28. Hillmann, F., Linde, J., Beckmann, N., Cyrulies, M., Strassburger, M., Heinekamp, T., Haas, H., Guthke, R., Kniemeyer, O., & Brakhage, A. A. (2014). The novel globin protein fungoglobin is involved in low oxygen adaptation of Aspergillus fumigatus. Molecular Microbiology, 93(3), 539– 553. https://doi.org/10.1111/mmi.12679

29. Kanj, A., Abdallah, N., & Soubani, A. O. (2018). The spectrum of pulmonary aspergillosis. In Respiratory Medicine (Vol. 141, pp. 121–131). W.B. Saunders Ltd. https://doi.org/10.1016/j.rmed.2018.06.029

30. Kousha, M., Tadi, R., & Soubani, A. O. (2011). Pulmonary aspergillosis: A clinical review. In European Respiratory Review (Vol. 20, Issue 121, pp. 156–174). European Respiratory Society. https://doi.org/10.1183/09059180.00001011

31. Kowalski, C. H., Kerkaert, J. D., Liu, K. W., Bond, M. C., Hartmann, R., Nadell, C. D., Stajich, J. E., & Cramer, R. A. (2019). Fungal biofilm morphology impacts hypoxia fitness and disease progression. Nature Microbiology, 4(12), 2430–2441. https://doi.org/10.1038/s41564-019-0558-7

32. Kowalski, C. H., Morelli, K. A., Schultz, D., Nadell, C. D., & Cramer, R. A. (2020). Fungal biofilm architecture produces hypoxic microenvironments that drive antifungal resistance. Proceedings of the National Academy of Sciences of the United States of America, 117(36), 22473–22483. https://doi.org/10.1073/pnas.2003700117

33. Kowalski, C. H., Morelli, K. A., Stajich, J. E., Nadell, C. D., & Cramer, R. A. (2021). A heterogeneously expressed gene family modulates the biofilm architecture and hypoxic growth of aspergillus fumigatus. MBio, 12(1), 1–21. https://doi.org/10.1128/mBio.03579-20

34. Lee, M. J., Geller, A. M., Bamford, N. C., Liu, H., Gravelat, F. N., Snarr, B. D., Le Mauff, F., Chabot, J., Ralph, B., Ostapska, H., Lehoux, M., Cerone, R. P., Baptista, S. D., Vinogradov, E., Stajich, J. E., Filler, S. G., Howell, P. L., & Sheppard, D. C. (2016). Deacetylation of fungal exopolysaccharide mediates adhesion and biofilm formation. MBio, 7(2), 1–14. https://doi.org/10.1128/mBio.00252-16

35. Lee, M. J., Liu, H., Barker, B. M., Snarr, B. D., Gravelat, F. N., Al Abdallah, Q., Gavino, C., Baistrocchi, S. R., Ostapska, H., Xiao, T., Ralph, B., Solis, N. V., Lehoux, M., Baptista, S. D., Thammahong, A., Cerone, R. P., Kaminskyj, S. G. W., Guiot, M. C., Latgé, J. P., … Sheppard, D. C. (2015). The Fungal Exopolysaccharide Galactosaminogalactan Mediates Virulence by Enhancing Resistance to Neutrophil Extracellular Traps. PLoS Pathogens, 11(10), e1005187. https://doi.org/10.1371/journal.ppat.1005187

36. Li, H., Barker, B. M., Grahl, N., Puttikamonkul, S., Bell, J. D., Craven, K. D., & Cramer, R. A. (2011). The Small GTPase RacA mediates intracellular reactive oxygen species production, polarized growth, and virulence in the human fungal pathogen aspergillus fumigatus. Eukaryotic Cell, 10(2), 174–186. https://doi.org/10.1128/EC.00288-10

37. Losada, L., Barker, B. M., Pakala, S., Pakala, S., Joardar, V., Zafar, N., Mounaud, S., Fedorova, N., Nierman, W. C., & Cramer, R. A. (2014). Large-Scale Transcriptional Response to Hypoxia in Aspergillus fumigatus Observed Using RNAseq Identifies a Novel Hypoxia Regulated ncRNA. Mycopathologia, 178(5–6), 331–339. https://doi.org/10.1007/s11046-014-9779-8

38. Lothier, J., Diab, H., Cukier, C., Limami, A. M., & Tcherkez, G. (2020). Metabolic responses to waterlogging differ between roots and shoots and reflect phloem transport alteration in medicago truncatula. Plants, 9(10), 1–20. https://doi.org/10.3390/plants9101373

39. Loussert, C., Schmitt, C., Prevost, M.-C. C., Balloy, V., Fadel, E., Philippe, B., Kauffmann-Lacroix, C., Latgé, J. P., & Beauvais, A. (2010). In vivo biofilm composition of Aspergillus fumigatus. Cellular Microbiology, 12(3), 405–410. https://doi.org/10.1111/j.1462-5822.2009.01409.x

40. Luo, W., & Brouwer, C. (2013). Pathview: An R/Bioconductor package for pathway-based data integration and visualization. Bioinformatics, 29(14), 1830–1831. https://doi.org/10.1093/bioinformatics/btt285

41. Maertens, J. A., Raad, I. I., Marr, K. A., Patterson, T. F., Kontoyiannis, D. P., Cornely, O. A., Bow, E. J., Rahav, G., Neofytos, D., Aoun, M., Baddley, J. W., Giladi, M., Heinz, W. J., Herbrecht, R., Hope, W., Karthaus, M., Lee, D. G., Lortholary, O., Morrison, V. A., … Ullmann, A. J. (2016). Isavuconazole versus voriconazole for primary treatment of invasive mould disease caused by Aspergillus and other filamentous fungi (SECURE): A phase 3, randomised-controlled, non-inferiority trial. The Lancet, 387(10020), 760–769. https://doi.org/10.1016/S0140-6736(15)01159-9

42. Maertens, J. A., Rahav, G., Lee, D. G., Ponce-de-León, A., Ramírez Sánchez, I. C., Klimko, N., Sonet, A., Haider, S., Diego Vélez, J., Raad, I., Koh, L. P., Karthaus, M., Zhou, J., Ben-Ami, R., Motyl, M. R., Han, S., Grandhi, A., & Waskin, H. (2021). Posaconazole versus voriconazole for primary treatment of invasive aspergillosis: a phase 3, randomised, controlled, non-inferiority trial. The Lancet, 397(10273), 499–509. https://doi.org/10.1016/S0140-6736(21)00219-1

43. Marr, K. A., Schlamm, H. T., Herbrecht, R., Rottinghaus, S. T., Bow, E. J., Cornely, O. A., Heinz, W. J., Jagannatha, S., Liang Koh, P., Kontoyiannis, D. P., Lee, D.-G., Nucci, M., Pappas, P. G., Slavin, M. A., Flavio Queiroz-Telles,;, Selleslag, D., Walsh, T. J., Wingard, J. R., & Maertens, J. A. (2015). Combination Antifungal Therapy for Invasive Aspergillosis A Randomized Trial. https://doi.org/10.7326/M13-2508

44. Moreno-Velásquez, S. D., Seidel, C., Juvvadi, P. R., Steinbach, W. J., & Read, N. D. (2017). Caspofungin-mediated growth inhibition and paradoxical growth in Aspergillus fumigatus involve fungicidal hyphal tip lysis coupled with regenerative intrahyphal growth and dynamic changes in β-1,3-glucan synthase localization. Antimicrobial Agents and Chemotherapy, 61(10). https://doi.org/10.1128/AAC.00710-17

45. Morino, Y., Kojima, H., & Tanase, S. (1979). Affinity labeling of alanine aminotransferase by 3-chloro-L-alanine. Journal of Biological Chemistry, 254(2), 279–285. https://doi.org/10.1016/s0021-9258(17)37915-2

46. Mowat, E., Lang, S., Williams, C., McCulloch, E., Jones, B., & Ramage, G. (2008). Phase-dependent antifungal activity against Aspergillus fumigatus developing multicellular filamentous biofilms. Journal of Antimicrobial Chemotherapy, 62(6), 1281–1284. https://doi.org/10.1093/jac/dkn402

47. Nierman, W. C., Pain, A., Anderson, M. J., Wortman, J. R., Kim, H. S., Arroyo, J., Berriman, M., Abe, K., Archer, D. B., Bermejo, C., Bennett, J., Bowyer, P., Chen, D., Collins, M., Coulsen, R., Davies, R., Dyer, P. S., Farman, M., Fedorova, N., … Denning, D. W. (2005). Genomic sequence of the pathogenic and allergenic filamentous fungus Aspergillus fumigatus. Nature, 438(7071), 1151– 1156. https://doi.org/10.1038/nature04332

48. Peña-Soler, E., Fernandez, F. J., López-Estepa, M., Garces, F., Richardson, A. J., Quintana, J. F., Rudd, K. E., Coll, M., & Cristina Vega, M. (2014). Structural analysis and mutant growth properties reveal distinctive enzymatic and cellular roles for the three major L-alanine transaminases of Escherichia coli. PLoS ONE, 9(7), e102139. https://doi.org/10.1371/journal.pone.0102139

49. Ries, L. N. A., Beattie, S., Cramer, R. A., & Goldman, G. H. (2018). Overview of carbon and nitrogen catabolite metabolism in the virulence of human pathogenic fungi. Molecular Microbiology, 107(3), 277–297. https://doi.org/10.1111/mmi.13887

50. Robert, V. A., & Casadevall, A. (2009). Vertebrate endothermy restricts most fungi as potential pathogens. Journal of Infectious Diseases, 200(10), 1623–1626. https://doi.org/10.1086/644642

51. Rocha, M., Licausi, F., Araújo, W. L., Nunes-Nesi, A., Sodek, L., Fernie, A. R., & van Dongen, J. T. (2010). Glycolysis and the tricarboxylic acid cycle are linked by alanine aminotransferase during hypoxia induced by waterlogging of Lotus japonicus. Plant Physiology, 152(3), 1501–1513. https://doi.org/10.1104/pp.109.150045

52. Sánchez, Ó. J., Ospina, D. A., & Montoya, S. (2017). Compost supplementation with nutrients and microorganisms in composting process. In Waste Management (Vol. 69, pp. 136–153). Elsevier Ltd. https://doi.org/10.1016/j.wasman.2017.08.012

53. Seidler, M. J., Salvenmoser, S., & Müller, F. M. C. (2008). Aspergillus fumigatus forms biofilms with reduced antifungal drug susceptibility on bronchial epithelial cells. Antimicrobial Agents and Chemotherapy, 52(11), 4130–4136. https://doi.org/10.1128/AAC.00234-08

54. Speth, C., Rambach, G., Lass-Flörl, C., Howell, P. L., & Sheppard, D. C. (2019). Galactosaminogalactan (GAG) and its multiple roles in Aspergillus pathogenesis. In Virulence (Vol. 10, Issue 1, pp. 976–983). Taylor and Francis Inc. https://doi.org/10.1080/21505594.2019.1568174

55. Surowiec, I., Karimpour, M., Gouveia-Figueira, S., Wu, J., Unosson, J., Bosson, J. A., Blomberg, A., Pourazar, J., Sandström, T., Behndig, A. F., Trygg, J., & Nording, M. L. (2016). Multi-platform metabolomics assays for human lung lavage fluids in an air pollution exposure study. Analytical and Bioanalytical Chemistry, 408(17), 4751–4764. https://doi.org/10.1007/s00216-016-9566-0

56. Taff, H. T., Mitchell, K. F., Edward, J. A., & Andes, D. R. (2013). Mechanisms of Candida biofilm drug resistance. In Future Microbiology (Vol. 8, Issue 10, pp. 1325–1337). Future Medicine Ltd London, UK. https://doi.org/10.2217/fmb.13.101

57. Wagener, J., & Loiko, V. (2018). Recent insights into the paradoxical effect of echinocandins. In Journal of Fungi (Vol. 4, Issue 1, p. 5). MDPI AG. https://doi.org/10.3390/jof4010005

58. Willger, S. D., Puttikamonkul, S., Kim, K.-H., Burritt, J. B., Grahl, N., Metzler, L. J., Barbuch, R., Bard, M., Lawrence, C. B., & Cramer, R. A. (2008). A Sterol-Regulatory Element Binding Protein Is Required for Cell Polarity, Hypoxia Adaptation, Azole Drug Resistance, and Virulence in Aspergillus fumigatus. PLoS Pathogens, 4(11), e1000200. https://doi.org/10.1371/journal.ppat.1000200

